# Context-dependent effects of developmental and adult diet on life-history traits in *Drosophila melanogaster*

**DOI:** 10.1101/2024.09.18.613663

**Authors:** Mohankumar Chandrakanth, Nishant Kumar, Chand Sura, Sudipta Tung

## Abstract

Life-history traits such as body size, reproduction, survival, and stress resistance are fundamental to an organism’s fitness and are highly influenced by nutritional environments across life stages. In this study, we employed a full factorial experimental design to investigate the effects of isocaloric diets (diets with equal caloric content but differing macronutrient composition) on key life-history traits in an outbred *Drosophila melanogaster* population. Our results demonstrated significant diet-induced plasticity, with male wing length (a proxy for body size) being influenced by the developmental diet; males reared on carbohydrate-rich developmental diets had larger wings as adults. Fertility increased with protein-rich diets at both developmental and adult stages, reaffirming the critical role of dietary protein in enhancing reproductive success. Lifespan exhibited sexually dimorphic responses to diet: carbohydrate-rich developmental diets extended male lifespan, while carbohydrate-rich adult diets reduced lifespan in both sexes. Stress resistance traits, including starvation and desiccation resistance, were unaffected by developmental diets but were influenced by adult diets, with carbohydrate-rich adult diets enhancing survival under both stress conditions in males and females. Importantly, while most traits exhibited additive effects of nutrition across life stages, a marginal interaction for male starvation resistance suggests that developmental and adult diets can interact in a trait- and sex-specific manner. Moreover, associations between dietary effects on life-history traits were context-dependent, driven primarily by adult diets and varying by sex. These findings emphasize the profound role of stage-specific nutritional environments in modulating life-history traits and their correlations, offering valuable insights into how organisms may adapt to changing ecological conditions and highlighting the importance of considering both developmental and adult dietary contexts in evolutionary studies.

## 1. Introduction

The nutritional environment directly impacts organismal fitness and, thereby, shapes their ecological and evolutionary trajectories by influencing various life history traits, including body size, reproductive output, longevity, and stress resistance (Flatt, 2020; Martin & Bize, 2018). Macronutrients like proteins, carbohydrates and lipids not only provide energy, but also serve as building blocks for protein, nucleotide, and lipid synthesis (Nehme et al., 2023; Prentice, 2005). They contribute to stored reserves like glycogen and lipids for future use (Olsen et al., 2021) and often act as signaling molecules that trigger growth, development, and cognitive functions (Rodriguez et al., 2017), thereby crucially influencing organismal life history and performance.

Several studies have investigated how dietary macronutrients influence life-history traits by using isocaloric diets – diets with identical caloric content but varying macronutrient composition – to disentangle macronutrient effects from those of total calories (Jang & Lee, 2018; Lee, 2015; Lee et al., 2008; Mair et al., 2005; Mudunuri et al., 2024) . Despite extensive research, the precise relationship between dietary macronutrients and life history traits remains unclear. For example, longevity, one of the most well-studied life history traits in the context of dietary manipulations, has shown diverse responses to varying dietary protein-to-carbohydrate (P:C) ratios in diets. Previous studies in *Drosophila* demonstrated that low P:C ratios (carbohydrate-rich) can have positive (Bruce et al., 2013; Lushchak et al., 2014) and complex non-monotonic (Lee, 2015) impacts on longevity. Similarly, while some studies have found that adult body size increases with high P:C ratios (Rodrigues et al., 2015), others have reported that body size maximizes at intermediate ratios (Lee & Roh, 2010). These observed variations are likely due to differences in the range of P:C ratios used in each study, highlighting the potential for varied outcomes.

Another potential reason for the observed differences in organismal performance across isocaloric dietary studies stems from the fact that all organisms undergo a pre-reproductive stage of growth and development, and food is crucial for both developmental and reproductive stages. However, dietary requirements during these two stages may differ significantly (Parks et al., 2020). Many previous studies have tested organismal performance by manipulating the diet at either the developmental (Dinh et al., 2022; Savola et al., 2022) or reproductive stage while keeping the diet constant for the developmental stage (Jang & Lee, 2018; Lee, 2015; Lee et al., 2008) or by applying the same dietary treatment to both stages (Mudunuri et al., 2024). While these approaches have provided valuable insights, they do not allow for the explicit examination of potential interactive effects between developmental and adult diets. This issue arises because it is not feasible to provide absolutely no food at one stage to study the impact of diet at the other stage on adult life history. Moreover, the interactive effects of stage-specific diet can vary across different life history traits (Klepsatel et al., 2020), further complicating the interpretation of such experimental data. In the simplest scenario, even if there is no significant interaction between the diets available across life stages, drastically different dietary requirements for developmental and reproductive stages (Al Shareefi & Cotter, 2019; Trumbo et al., 2002) can still generate a non-monotonic relationship. Therefore, the observed non-monotonic diet-induced plasticity profiles of various life-history traits likely reflect the consequences of mismatched stage-specific dietary needs. Additionally, interactive effects between diets at different life stages, or a combination of both factors could contribute to these patterns. One approach to delineate these effects involves a full factorial experimental design incorporating stage-specific diet availability, but such designs are less commonly used due to logistical challenges (see, however, Duxbury & Chapman, 2020; Poças et al., 2022; Pullock et al., 2023; Ruchitha et al., 2024).

Additionally, investigating the effects of diet on multiple life-history traits under diverse dietary environments offers an opportunity to explore how dietary changes influence these traits in a coordinated manner. Associations between dietary effects on life-history traits arise because diets impact the allocation of finite energy resources in an organism, shaping multiple traits simultaneously rather than in isolation. For example, dietary interventions often reveal trade-offs between energy-intensive traits such as reproductive output, longevity, and stress resistance, reflecting the energetic constraints organisms face (Fanson & Taylor, 2012; Le Rohellec & Le Bourg, 2009; Lee et al., 2008; Zanco et al., 2021). Similarly, body size, which may indirectly reflect stored resources such as glycogen and lipids, is influenced by both dietary composition and caloric content (Poças et al., 2022). Across individuals, larger body size is often positively associated with fecundity (Honek, 1993), longevity (Tantawy & Vetukhiv, 1960) and stress resistance (Harshman et al., 1999). Together, these findings highlight the importance of understanding how dietary effects on one trait may cascade to influence other traits, offering critical insights into the evolutionary and ecological consequences of diet-induced plasticity. Moreover, such insights have translational implications for conservation, agriculture, and health sciences, offering strategies to improve health and longevity, manage wildlife and pest populations, and optimize breeding programs for livestock and economically important insects. (Baenas & Wagner, 2019; Ruden et al., 2005; Solovev et al., 2018; Vijendravarma et al., 2012)

While significant progress has been made in understanding the effects of diet on individual life-history traits, the associations between dietary effects on multiple traits remain relatively underexplored. Several studies have provided valuable insights into specific traits or limited subsets of traits (Klepsatel et al., 2020; Krittika & Yadav, 2022; May et al., 2015; Mirth et al., 2019). However, comparing findings across studies can be challenging due to variations in experimental designs, such as differences in genetic backgrounds, diet composition, caloric content, and the use of solid versus liquid food (Gillette et al., 2020; Lüersen et al., 2019). Building on this body of work, there is a need for an experimental framework to examine key fitness-related traits—including longevity, reproductive output, and stress resistance—under common isocaloric dietary regimes with contrasting macronutrient compositions.

We addressed the existing knowledge gaps using a full-factorial experimental design with isocaloric diets that varied in macronutrient composition, specifically the proportions of proteins and carbohydrates. By isolating macronutrient effects from overall caloric intake, our approach allowed us to dissect the impacts of diet composition during developmental and adult stages. Using *Drosophila melanogaster* as a model system, we leveraged its highly conserved nutrition-responsive pathways and distinct larval (developmental) and adult feeding stages to investigate the dietary effects across life stages. In the wild, *Drosophila* inhabits ephemeral food sources such as ripening and decaying fruits, where macronutrient composition changes dynamically over time. These natural dietary fluctuations expose flies to varying protein-to-carbohydrate (P:C) ratios across their feeding stages, providing a relevant ecological and evolutionary context for investigating nutrient-dependent trade-offs and their fitness consequences (Markow & O’Grady, 2008; Shu et al., 2022). Our laboratory-based approach complemented this natural ecological context by enabling precise manipulation of dietary conditions to isolate the effects of developmental and adult diets, while minimizing confounding factors which are difficult to control in natural settings. To ensure our findings were robust and generalizable, we used a large, outbred population of *Drosophila melanogaster*, representing large standing genetic diversity similar to natural populations.

Across the experimental dietary regimes, we examined five key life-history traits: wing length, fertility, longevity, desiccation resistance, and starvation resistance. Wing length was used as an indirect proxy for body size, which in turn is often correlated with stored energetic resources such as lipids and glycogen in insects (Enriquez et al., 2022; Güler et al., 2015; Shin et al., 2012). These reserves influence how individuals allocate energy to life-history traits such as reproduction, survival, and stress resistance (Chippindale et al., 1996; Harshman & Zera, 2007; Zera & Harshman, 2001). Fertility and longevity were included as primary fitness-related traits due to their direct contributions to organismal fitness and their high energy demands (Flatt, 2020). Desiccation and starvation resistance were selected as representative stress-resistance traits, particularly relevant in natural contexts where resource availability fluctuates (Marron et al., 2003). Collectively, these traits provide a comprehensive framework to investigate resource allocation, trade-offs, and fitness strategies in response to stage-specific diet manipulations. Moreover, by comparing associations between the dietary effects on these traits, we explored how macronutrient composition at different life stages influences fitness in a sex-specific manner.

Our results demonstrated largely additive effects of developmental and adult diets in shaping life-history traits, though a marginal interaction for male starvation resistance suggests potential trait- and sex-specific dietary interactions. Additionally, we highlight stage- and sex-specific dietary effects and show that associations between these dietary effects on the life history trait are context-dependent. By leveraging stage-specific dietary macronutrient manipulations in an isocaloric framework, this work advances our understanding of life-history consequences and provides novel insights into how nutritional environments influence diet-induced plasticity and their impacts on evolutionary dynamics.

## 2. Materials and Methods

### 2.1 Ancestry and maintenance of the experimental Population

The experimental population used in this study traces its ancestry to the large, outbred, laboratory-maintained *Drosophila melanogaster* Baseline Population (BP) (Mudunuri et al., 2024). The BP population was established by equally mixing four Melanogaster Baseline (MB_1–4_) populations, which originated from a wild-caught IV population collected in South Amherst, MA, USA, in 1970 (Ives, 1970) and have since been maintained in the laboratory as large, outbred populations (Sarangi et al., 2016; Shenoi et al., 2016).

The BP population was reared for 30 generations on corn-sugar-yeast medium (CSY) with protein-to-carbohydrate (P:C) ratio of 0.4 having caloric content of 714 kcal/L, hereafter referred to as the baseline medium (see Table S1 for detailed recipe).

The baseline populations, was maintained on a 14-day discrete generation cycle under standard laboratory conditions (25°C, >60% relative humidity, and constant light as per the ancestral maintenance regime). On the 13^th^ day post-egg collection, an egg-laying substrate of baseline medium was provided, and flies were allowed to lay eggs overnight for approximately 14 hours to initiate the next generation. Eggs were carefully scraped and washed in double-distilled water using a fine paintbrush and aliquoted ∼300-350 eggs into per plastic fly-culturing bottle (Laxbro® FLBT-20) containing 50 ml baseline medium (see Text S1 for egg collection protocol). Seven bottles were used per population, yielding a total population size of ∼2100 flies. On the 11^th^ day post-egg collection, all eclosed adult flies were transferred to a plexiglass cage (25 cm × 20 cm × 15 cm) and provided with fresh food every other day until egg collection for the next generation.

### 2.2 Experimental diet regimes and generation of the experimental flies

In this study, we took two isocaloric diets with distinct compositions: a carbohydrate-rich diet with a P:C ratio of 0.25 (henceforth referred to as the C diet) and a protein-rich diet with a P:C ratio of 0.7 (henceforth referred to as the P diet). Both diets had an equivalent caloric density of 714 kcal/L and were formulated following established protocols (Mudunuri et al., 2024; see Table S1 for details). Both of the P and C diets were provided during the larval and adult stages in a full factorial design, resulting in four distinct experimental diet regimes: CC, CP, PC, and PP. The first letter indicates the larval diet, and the second letter indicates the adult diet. For example, CP represents a diet regime that begins with a carbohydrate-rich larval diet followed by a protein-rich adult diet. Flies were exposed to their assigned diets in a no-choice setup during both developmental and adult stages. While diets were isocaloric by design, actual intake or assimilation was not measured; dedicated assays would be needed to quantify these parameters.

Experimental flies were generated by aliquoting approximately 150 eggs into each fly-culturing bottle containing 50 ml of the designated larval diet. This ensured consistent rearing conditions by minimizing larval crowding. The number of bottles was adjusted based on the total number of adult flies required for each specific assay.

To enable the larval-to-adult diet transition, we customized fly-culturing bottles with removable bottoms (see Figure S1). At the late pupal stage, the larval food base was replaced with the designated adult food. This ensured adults received the appropriate diet immediately upon eclosion. To account for the developmental delays associated with the C larval diet (Mudunuri et al., 2024) , the diet switch was performed on day 7 post-egg collection for PC and PP regimes and on day 8 for CC and CP regimes.

Flies reared under these diet regimes were then assayed for five key adult life-history traits on the 12^th^ day post-egg collection, with all adults fully eclosed and having consumed respective larval and adult diets prior to the experiments.

### 2.3 Wing length assay

To assess the effects of stage-specific diet composition on body size, we measured wing length as a proxy in both males and females, following established methods (Joubert & Bijlsma, 2010; Mudunuri et al., 2024). One wing from each fly was randomly selected and clipped using fine forceps. The clipped wings were mounted on a glass slide using PEG 6000 (SRL, Cat: 25322-68-3). The images of mounted wings were captured using a Zeiss Stemi 305 microscope with Axiocam 105 colour camera under 40X magnification. The final measurement of the wing was done by measuring the length between anterior cross vein (ACV) to L3 nodes (Güler et al., 2015; Joubert & Bijlsma, 2010) using ImageJ software, calibrated with a standard scale (Stage micrometer-ESM11).

The assay included 30 replicates per sex for each of the four diet regimes, resulting in measurements from 240 wings (30 replicates × 2 sexes × 4 regimes). Although we did not explicitly account for directional asymmetry in *Drosophila* wings (Møller & Thornhill, 1998), random selection of a single wing per individual likely minimized systematic bias. Future studies could improve accuracy by explicitly considering directional asymmetry in wing measurements.

### 2.4 Fertility assay

In order to estimate the reproductive output across experimental diet regimes, we collected mated flies on the 12^th^ day post-egg collection for assay setup. Flies were sorted by sex under brief CO_2_ anaesthesia (less than 1 minute) and two males and two females were randomly placed in a glass vial containing the respective adult diet. A total of 30 such replicates were created for each diet regime. These vials were incubated for 24 hours under standard laboratory conditions to allow egg laying. Afterward, adult flies were discarded, and the vials containing eggs were incubated for eggs to develop into adults. Based on a previous study using flies from the same genetic background, which demonstrated no differential pre-adult survivorship between the experimental P and C diets (Mudunuri et al., 2024) the number of eclosed adult offsprings was recorded as the measure of fertility for each vial. Fertility per mating pair was calculated and subjected to statistical analysis. This fertility assay involved a total of 480 flies for all four regimes (30 replicates × 2 flies × 2 sex × 4 regimes).

### 2.5 Longevity assay

For measuring lifespan of flies across four experimental diet regimes, 8 adult males and 8 adult females were separately sorted under brief CO_2_ anaesthesia (less than 1 minute) into vials containing ∼5ml of respective adult diet. Ten replicates were prepared for each sex in each diet regime, and incubated under standard laboratory conditions. Each vial was monitored daily to record the number of alive flies per vial. Fresh food was provided every third day until all flies in the vial had died. The lifespan of flies was estimated as the number of days from the assay setup to the death of each fly. This setup assessed lifespan of 640 flies in total (10 replicates × 8 flies × 2 sex × 4 regimes). Individual death data were analysed statistically, with vial replicate as a random factor.

### 2.6 Starvation resistance assay

To assess starvation resistance, 8 males and 8 females were placed in separate vials for each replicate, with 10 replicates per sex per diet regime. Each vial contained ∼3mL 1% agar and was incubated under standard laboratory conditions. Vials were monitored every 4 hours for counting the number of alive flies per vial, and fresh agar was provided every second day until all flies in a vial had died. Starvation resistance was measured as the number of hours from the assay setup to the death of each fly. This setup assayed 640 flies (10 replicates × 8 flies × 2 sex × 4 regimes) across all four regimes. Individual mortality data were analysed statistically, with vial identity included as a random factor.

### 2.7 Desiccation resistance assay

Desiccation resistance across experimental diet regimes was determined by sorting eight males and eight females into separate empty vials without any source of food or water. 10 vial-replicates were prepared for each diet regime × sex combination. Vials were monitored every two hours to count the number of dead flies until all flies in the vial had died.

Desiccation resistance was recorded as the number of hours each fly survived after the assay began. In total, this assay measured the desiccation resistance of 640 flies (10 replicates × 8 flies × 2 sex × 4 regimes) across the four regimes. Individual mortality data were analysed statistically, with the vial identity as random factor.

### 2.8 Statistical analysis

To evaluate the effects of larval and adult diets on wing length, lifespan, starvation resistance, and desiccation resistance, we first fitted a full factorial model including larval diet, adult diet, and sex as fixed factors. All models were constructed based on the full factorial experimental design, and no stepwise simplification was applied, as all fixed effects and interactions were pre-specified based on biological hypotheses. For lifespan, starvation resistance, and desiccation resistance, vial identity was included as a random factor. We observed a significant main effect of sex across all these traits, and significant three-way interactions (larval diet × adult diet × sex) for lifespan and starvation resistance (Supplementary Table S2). Given these results and the well-established sexual dimorphism of wing length, lifespan, starvation resistance, and desiccation resistance in *Drosophila*, we performed sex-stratified analyses for these traits to ensure consistency and enable clear interpretation of sex-specific dietary effects.

Wing length data for each sex were analysed using linear regression with larval and adult diet as fixed factors. We assessed model assumptions using residual diagnostics. Q-Q plots showed that residuals closely aligned with a normal distribution, and residuals vs. fitted plots showed no major heteroscedasticity or non-linear trends (Figure S2 for females, Figure S3 for males), confirming that model assumptions were adequately met.

Fertility data, expressed as count of viable offspring, exhibited overdispersion when modelled using a Poisson distribution (dispersion ratio = 2.08, Pearson’s χ² = 241.79, p < 0.001; *performance::check_overdispersion*). Therefore, we fitted a generalized linear model with a negative binomial distribution and log link (*MASS::glm.nb*) including larval diet and adult diet as fixed factors, which eliminated overdispersion (dispersion ratio = 0.93, p = 0.664) and provided a substantially improved fit (reduction in residual deviance = 131.15).

For lifespan, starvation resistance, and desiccation resistance, survivorship curves were analysed using a Cox proportional hazards model. Survival objects were created with the *Surv* function from the *survival* package, and a mixed-effects Cox model was fitted with the *coxme* function from the *coxme* package. The model included developmental diet, adult diet, and their interaction as fixed factors, with vial identity treated as a random factor. For these three traits, proportional hazards assumptions were assessed using log-minus-log survival plots generated with *survival::survfit()* and *survminer::ggsurvplot(fun = “cloglog”)* in R. Curves stratified by larval and adult diet were evaluated separately for each sex (Figure S4-S9), with approximate parallelism interpreted as support for the proportional hazards assumption. The sex-stratified analyses results tables for all these traits are provided in the supplementary material (Table S3 – Table S7).

Model fit was assessed using the *r2* function from the *performance* package in R. For linear models (e.g., wing length), we report adjusted R^2^. For count data (e.g., fertility, modeled using negative binomial regression) and survival data (e.g., lifespan, starvation, and desiccation resistance, modelled using Cox proportional hazards models as this analysis is not supported for mixed-effects Cox models), we report Nagelkerke’s R^2^. These metrics reflect the proportion of variance explained by fixed effects and are summarized in Supplementary Table S8. Given the complexity and multifactorial nature of life-history traits, and the use of genetically variable outbred populations, relatively low R^2^ values are expected and consistent with previous literature (Houle, 1992; Nakagawa & Schielzeth, 2013)

To complement our statistical analysis, we also report descriptive statistics and effect size estimates (Cohen’s *d*; J. Cohen, 1988) for all treatment comparisons in Supplementary Table S9, interpreted as very small (*d* < 2), small (2 ≤ *d* < 5), medium (5 ≤ *d* < 8), and large (*d* ≥ 8). Visualizations of the effects of developmental and adult diets on wing length and fertility were generated using the *ggplot2* package. Survival curves for lifespan, starvation resistance, and desiccation resistance were plotted using the *survminer* and *patchwork* packages. All analyses and visualizations were conducted in R version 4.2.1 and RStudio v2022.06.23 (RStudio Team, 2020).

### 2.9 Dietary-effect Index

To analyze the pairwise associations between dietary effects on measured traits, we conducted separate analyses for each combination of developmental stage (developmental or adult diet) and sex (males and females). For each combination, we first calculated the log response ratio for each trait (lnRR_trait_). The quantifies proportional changes in mean trait values between protein-rich (P) and carbohydrate-rich (C) diets at each stage: lnRR_trait_ = ln(X_P_/X_c_), where X_P_ and X_c_ are the mean trait values under the P and C diets, respectively. A positive lnRR_trait_ indicates the trait increased under the P diet, while a negative value indicates a decrease. The sampling variance of lnRR_trait_ for each trait was calculated as: V_trait_ = (S_P_ / X_P_)^2^ / N_P_ + (S_C_ / X_C_)^2^ / N_C_, where S_P_ and S_C_ are the standard deviations of the trait, and N_P_ and N_C_ are the sample sizes under the P and C diets, respectively.

To assess the direction of association between dietary effect on two traits, we computed the Dietary-effect Index (DEI*_ij_*) for each pair of traits (*i* and *j*) by taking the ratio of the their lnRR_trait_ values: DEI*_ij_* = lnRR_trait*i*_ / lnRR_trait*j.*_ Assuming dietary effects on the two traits are independent, the standard error of DEI*_ij_* was estimated as: SE*_ij_* ≈ √ (V_trait*i*_ /(lnRR_trait*j*_)^2^ + (lnRR_trait*i*_)^2^×V_trait*j*_/(lnRR_trait*j*_)^4^) (van Kempen & van Vliet, 2000).We then computed the z-score to evaluate the statistical significance of the direction of association: z = DEI*_ij_*/SE*_ij_*Under the null hypothesis (DEI*_ij_* = 0), there is no meaningful directional association between the dietary effects on the two traits, implying that the changes in the two traits are unrelated or randomly aligned. Positive or negative DEI*_ij_* values indicate the direction of the association. The corresponding p-value was used to assess statistical significance. For example, a significantly positive DEI*_ij_* for starvation resistance and desiccation resistance under adult diet suggests that this stage-specific diet not only increases starvation resistance but also enhances desiccation resistance. Conversely, a negative DEI*_ij_* for fertility and desiccation resistance under the same diet implies a trade-off, where higher reproductive output is associated with reduced desiccation resistance, and vice versa.

## 3. Results

The full factorial analysis demonstrated significant main effects of sex for wing length (F = 528.2, p < 0.001), lifespan (χ² = 79.2, p < 0.001), starvation resistance (χ² = 66.1, p < 0.001), and desiccation resistance (χ² = 159.4, p < 0.001). Most importantly, we detected significant three-way interactions (larval diet × adult diet × sex) for lifespan (χ² = 6.6, p = 0.010) and starvation resistance (χ² = 5.3, p = 0.022), indicating that optimal dietary combinations are sex-specific (Table S2). The sex-stratified analyses presented below showed these distinct nutritional requirements (Tables S3–S7), with model fit metrices and effect sizes (Cohen’s *d*) provided in Tables S8–S9.

### 3.1 Developmental diet composition primarily affects wing length in male flies

We found that the impact of diet on wing length was developmental stage and sex specific. There was no significant effect of developmental diet on female wing length (Figure 1A; F =1.9, *p* = 0.17). In contrast, male flies provided with carbohydrate-rich larval diet had greater average wing length compared to those provided with protein-rich larval diet (Figure 1B; F = 6.8, *p* = 0.01). The effect size of this difference was small (Cohen’s *d* = 0.35; Table S9). The composition of the adult diet had no significant effect on wing length in either sex (Figure S2A; F =0.2, *p* = 0.62 for females and Figure S2B; F =0.2, *p* = 0.59 for males) and no significant interaction was observed between larval diet and adult diet in either of the sexes (Table S3; F =1.8, *p* = 0.18 for females, and F =0.1 *p* = 0.73 for males).

**Figure 1.**
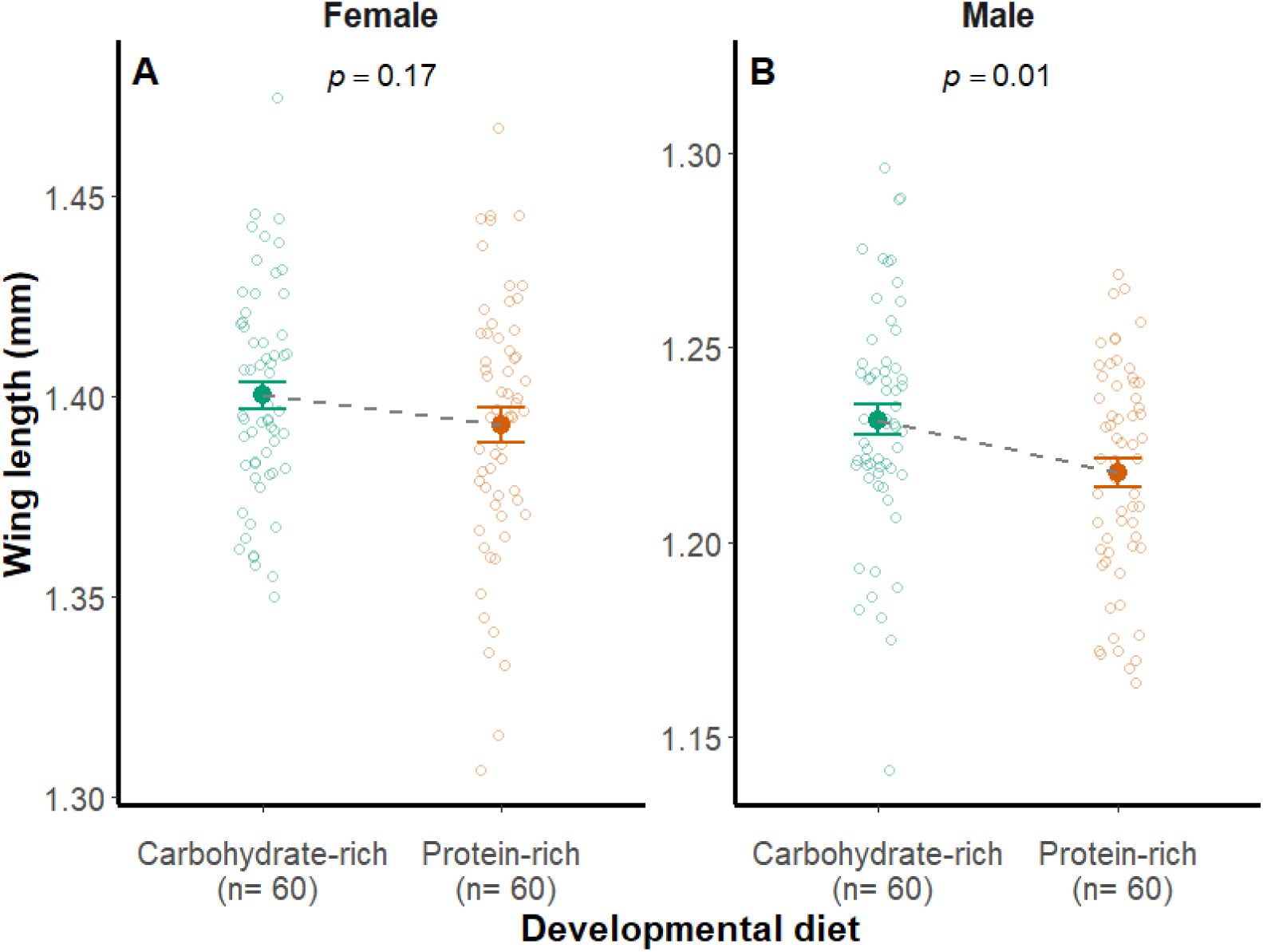
Effect of developmental diet composition on wing length. The graphs depict the impact of carbohydrate-rich (C) and protein-rich (P) developmental diet on adult wing length (a proxy for body size) of female (A) and male (B) flies, respectively. The error bar depicts the standard error (± SE) of mean for each diet provided at developmental stage. The filled circles indicate mean wing length, while small open circles denote wing length of individual flies in each group. Teal green and orange-red color represents the carbohydrate-rich and protein-rich isocaloric diets, respectively. n denotes the sample size. *p*-values indicate the statistical significance of dietary effects, estimated using a generalized linear model with a Gaussian distribution (GLM, identity link). See Supplementary Table S3 for full statistical outcome.

### 3.2 Fertility is influenced by both developmental and adult diets

Flies raised on a protein-rich developmental diet produced more viable offspring compared to those on a carbohydrate-rich developmental diet (Figure 2A, F = 4.38, *p* = 0.04). The effect size of this difference was small (Cohen’s *d* = 0.31; Table S9). Adult diet had a particularly strong effect on fertility (Figure 2B, F = 61.5, *p* < 0.001), with flies provided with protein-rich adult diet producing more offspring than those fed a carbohydrate-rich adult diet and the effect size of difference was large (Cohen’s *d* = 1.41; Table S9). The interaction between developmental diet and adult diet was not significant (Table S4, F= 0.42, *p* = 0.5).

**Figure 2.**
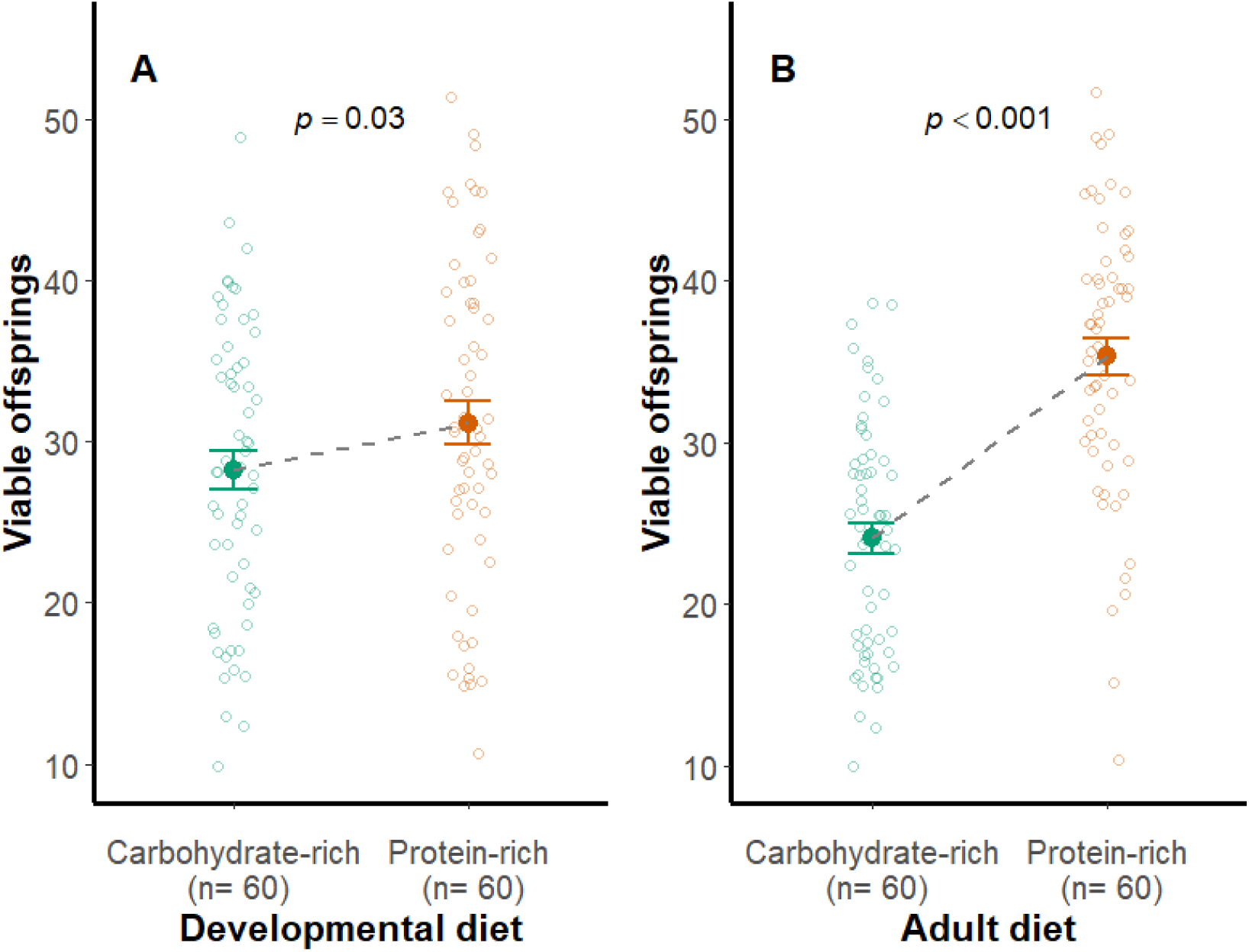
Impact of both developmental diet and adult diet on the number of viable offsprings. The graphs depict the impact of carbohydrate-rich (C) and protein-rich (P) developmental diet (A) and adult diet (B) on fertility, measured as the number of viable offsprings produced. The error bar depicts the standard error (± SE) of mean and the filled circles indicate mean wing length, while small open circles denote wing length of individual flies in each group. Teal green and orange-red color represents the carbohydrate-rich and protein-rich isocaloric diets. n represents the sample size. *p*-values indicate the statistical significance of dietary effects derived from a generalized linear model with a negative binomial distribution. See Supplementary Table S4 for full statistical outcome.

### 3.3 Longevity is affected by developmental and adult diets in sexually dimorphic manner

We observed that the dietary effects on lifespan vary by developmental stage and sex. In females, adult diet was the primary determinant of lifespan, with protein-rich adult diet extending mean lifespan by approximately 34% (37.3 vs 27.8 days; Figure 3B; Table S5, χ² = 35.8, p < 0.001, Cohen’s *d* = 0.89; Table S9). Developmental diet had no significant effect on female longevity (Figure 3A; Table S5, χ² = 0.1, p = 0.75).

**Figure 3.**
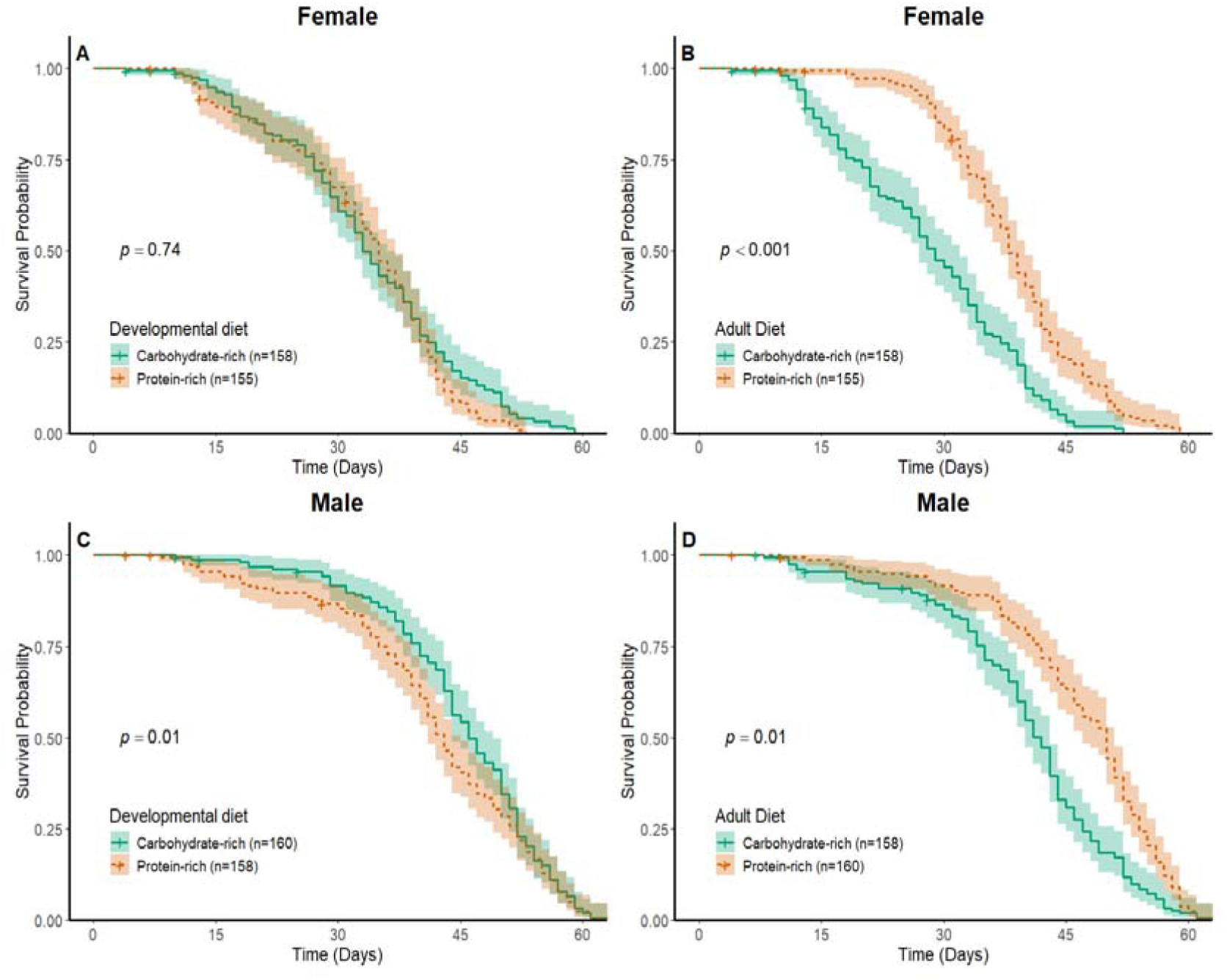
Impact of developmental diet and adult diet on survival probability in female and male flies. The survival curves illustrate the impact of carbohydrate-rich (C, teal green) and protein-rich (P, orange-red) diets provided during larval (A and C) and adult (B and D) stages on survival probability. Panels (A) and (B) show the effects of diets on females, while panels (C) and (D) display the effects of diets on male flies. The shaded area around the curve represents 95% confidence interval. Teal green and orange-red color represents the carbohydrate-rich and protein-rich isocaloric diets. n represents the sample size. *p*-values indicate the statistical significance of dietary effects assessed via a Cox proportional hazards mixed-effects model. See Supplementary Table S5 for full statistical outcome.

In contrast, male longevity was significantly influenced by developmental diet (Figure 3C, χ^2^=6.2, *p=* 0.01), with a carbohydrate-rich developmental diet extending adult lifespan (Small effect size, Cohen’s *d* = 0.20; Table S9). Similar to females, the adult diet impacted male lifespan (Figure 3D, χ^2^= 6.5, *p* = 0.01), with flies fed a protein-rich adult diet living longer than those fed on a carbohydrate-rich adult diet (Medium effect size, Cohen’s *d* = 0.51; Table S9). The interaction between developmental diet and adult diet was not significant for either sex (Table S5; χ^2^= 2.8, *p* = 0.09 for males and χ^2^=2.4, *p* =0.11 for females).

### 3.4 Stage-specific dietary effects on starvation resistance differ between sexes

Starvation resistance in females was not significantly influenced by developmental diet (Figure 4A; χ^2^ = 0.9, *p* = 0.33). However, adult diet had a marginally significant effect (Figure 4B; χ^2^ = 3.9, *p* = 0.046), with females reared on carbohydrate-rich adult diet surviving longer under starvation than those on protein-rich adult diet (Small effect size, Cohen’s *d* = 0.39; Table S9). There was no significant interaction between developmental and adult diets (χ^2^ = 0.5, *p* = 0.47). Whereas, in males, starvation resistance showed a more complex response to stage-specific diet composition with a marginally significant interaction between developmental and adult diets (Figure 4C–D; Table S6; χ^2^ = 3.6, *p* = 0.05).

**Figure 4.**
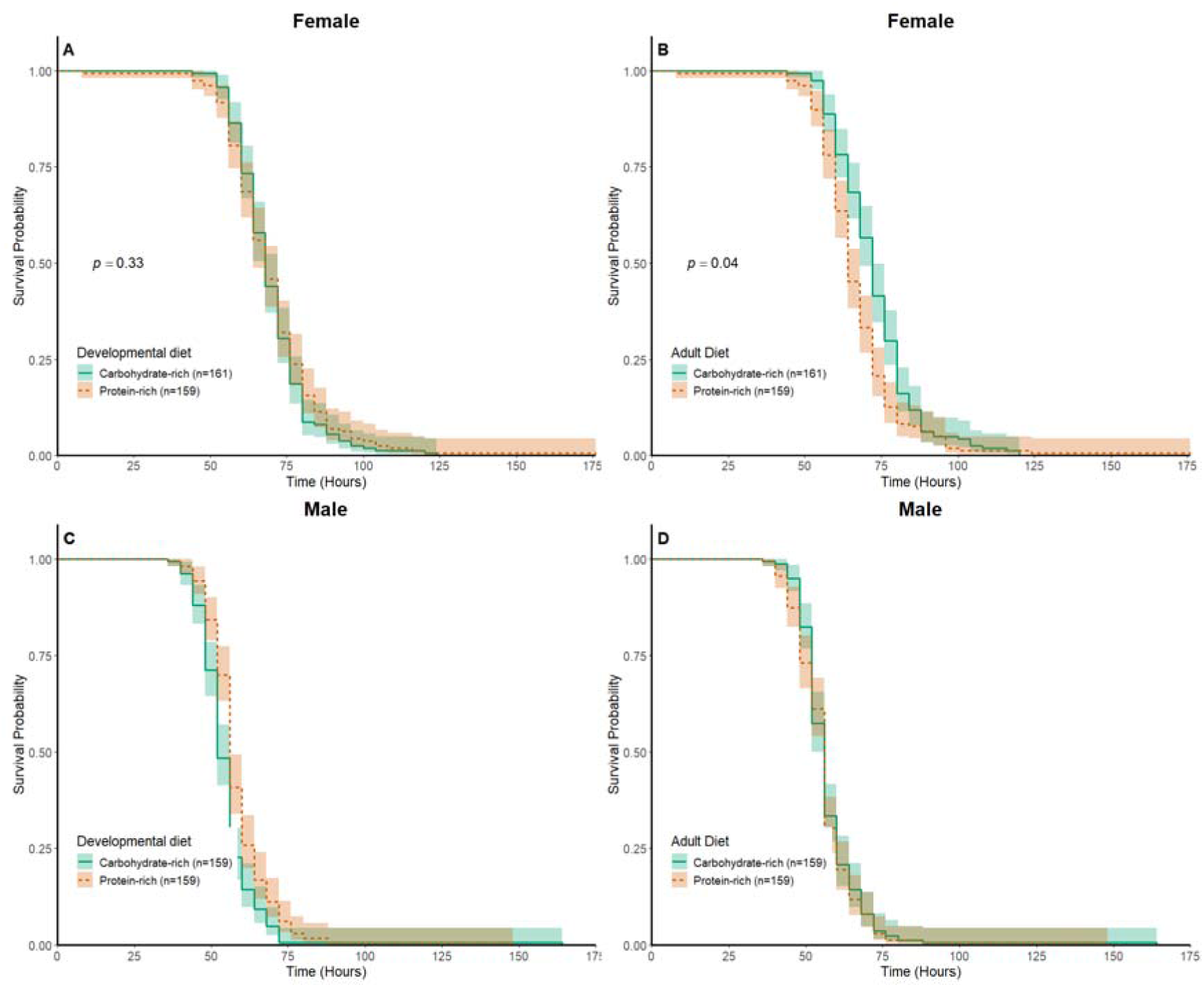
Impact of developmental diet and adult diet on survival probability under starvation stress in female and male flies. The survival curves depict the influence of carbohydrate-rich (C, teal green) and protein-rich (P, orange-red) diets provided during larval (A and C) and adult (B and D) stages on survival probability under starvation stress. Panels (A) and (B) represents the effects of diets on female flies, while panels (C) and (D) display the effects of diets on male flies. The shaded regions around the curve represents 95% confidence interval. Teal green and orange-red color represents the carbohydrate-rich and protein-rich isocaloric diets. n denotes the sample size. *p*-values indicate the statistical significance of dietary effects assessed via a Cox proportional hazards mixed-effects model. Absence of *p*-values denotes a significant interaction between developmental and adult diets. See Supplementary Table S6 for full statistical outcome.

### 3.5 Desiccation resistance of flies is primarily affected by diet provided during adult stage

Developmental diet did not affect desiccation resistance in either females (Figure 5A, χ^2^= 0.5, *p* = 0.46) or males (Figure 5C, χ^2^= 2.9, *p* = 0.08). In contrast, the adult diet composition had a significant impact in both females (Figure 5B, χ^2^= 7.3, *p* = 0.006; Table S9, Cohen’s *d* = 0.4) and males (Figure 5D, χ^2^= 11.1, *p* = <0.001; Table S9, Cohen’s *d* = 0.58), with carbohydrate-rich adult diet leading to greater desiccation resistance in both the sexes. The interaction between developmental diet and adult diet on desiccation resistance was not significant in either sex (Table S7, χ^2^= 0.01, *p* = 0.89 in females and χ^2^ <0.0001, *p* = 0.99 in males).

**Figure 5.**
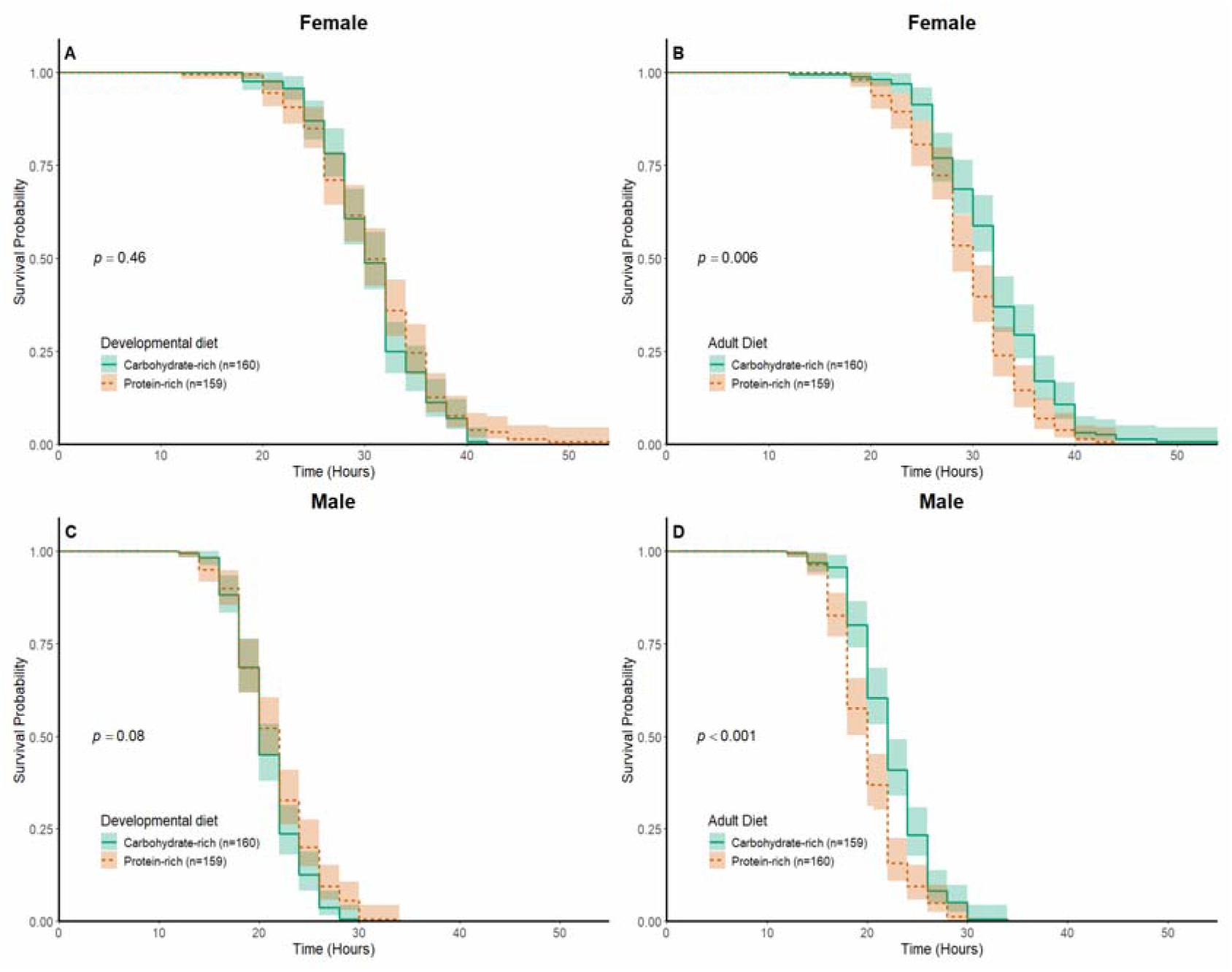
Impact of developmental diet and adult diet on survival probability under desiccation stress in female and male flies. The survival curves illustrate the effect of carbohydrate-rich (C, teal green) and protein-rich (P, orange-red) diets provided during larval (A and C) and adult (B and D) stages on survival probability under desiccation stress. Panels (A) and (B) show the effects of diets on female flies, while panels (C) and (D) display the effects of diets on male flies. The shaded area around the curve represents 95% confidence interval. Teal green and orange-red color represents the carbohydrate-rich and protein-rich isocaloric diets. n represents the sample size. *p*-values indicate the statistical significance of dietary effects assessed via a Cox proportional hazards mixed-effects model. See Supplementary Table S7 for full statistical outcome.

### 3.6 Context dependent association of dietary effects on life-history traits

We found no significant associations between the effects of developmental diet on the measured life-history traits in either males or females (Table S10). However, for the effects of adult diet, we identified sex-specific associations involving fertility, starvation resistance, longevity, and desiccation resistance (Table 1, Table S11). In females, adult diet effects on starvation resistance were negatively associated with both fertility and longevity, while positively associated with desiccation resistance. In contrast, these associations were not significant in males. Additionally, in both sexes, the effects of adult diet on fertility and longevity were positively associated, but these effects were negatively associated with dietary effects on desiccation resistance. Detailed statistics and calculations are provided in the Supplementary file Table S10-S11.

**Table 1.** Pairwise associations between adult dietary effects on life-history traits in females (upper triangle) and males (lower triangle). Methodological details are provided in Section 2.9; see Supplementary Tables S10 and S11 for statistical formulae and test outputs. *p*-values indicate the statistical significance of associations of dietary effects on trait combinations.

| ♀<br>♂ | Wing length | Fertility | Longevity | Starvation resistance | Desiccation resistance |
| --- | --- | --- | --- | --- | --- |
| Wing length | – | Non-significant | Non-significant | Non-significant | Non-significant |
| Fertility | Non-significant | – | Positive<br>($p = 7.12\text{e-}08$) | Negative<br>($p = 0.001$) | Negative<br>($p = 0.001$) |
| Longevity | Non-significant | Positive<br>($p = 0.0001$) | – | Negative<br>($p = 0.002$) | Negative<br>($p = 0.001$) |
| Starvation resistance | Non-significant | Non-significant | Non-significant | – | Positive<br>($p = 0.012$) |
| Desiccation resistance | Non-significant | Negative<br>($p = 0.00002$) | Negative<br>($p = 0.0006$) | Non-significant | – |

## 4. Discussion

Life-history traits are deeply intertwined with organismal response to the environmental cues, including the availability and composition of nutrients at various life stages. In this study, we investigated the effects of dietary composition by altering the relative proportion of proteins and carbohydrates in both developmental and adult stages in *Drosophila melanogaster*. Using a full factorial design, we assessed the effects of these stage-specific dietary manipulations on key life-history traits-wing length, fertility, longevity, starvation resistance and desiccation resistance-and evaluated pair-wise associations between dietary effects on these traits. Our results showed significant effect of both developmental and adult diet on these traits, with largely no significant interaction between the two stages. Additionally, we observed sexually dimorphic dietary impacts in some cases. Pairwise associations between dietary effects on life-history traits were found to be context-dependent, varying by life stage and sex.

Body size is a key determinant of an organism’s fitness, influencing its various aspects such as mating success, lifespan, avoiding predation, and stress resistance (J. E. Cohen et al., 1993; Rose & Mueller, 1993) . In our study, wing length (a proxy for body size) was influenced by diet in a sex- and stage-specific manner. Male wing length was significantly affected by the developmental diet, with carbohydrate-rich diets resulting in slightly larger wings compared to protein-rich diets. In contrast, neither developmental nor adult diets influenced wing length in females. These findings align with previous studies emphasizing the predominant role of juvenile or developmental diets in determining adult body size (Callier & Nijhout, 2013; Nijhout, 2003; Nijhout et al., 2014) , including evidence that appendage size, such as wing area and femur length, is primarily shaped by developmental diet in *Drosophila melanogaster* (Poças et al., 2022). This highlights the critical role of developmental nutrition in influencing adult body size, with the observed male-specific effect suggesting potential sex-based differences in resource allocation during development. One plausible explanation is that males reared on carbohydrate-rich diets compensate for protein limitation by consuming more food (Almeida de Carvalho & Mirth, 2017), leading to increased fat storage, which has been positively associated with larger pupal size and, consequently, bigger adult body size in *D. melanogaster* (Enriquez et al., 2022). Additionally, we found no interaction between developmental and adult diets for this trait, suggesting that the effects of stage-specific diets on wing length are independent. Together, these results highlight the nuanced nature of diet-induced plasticity in body size, indicating sex-specific differences in resource allocation and providing a foundation to explore its implications for other energy intensive life-history traits such as reproductive output, lifespan and stress resistance traits, which we discussed below.

Reproduction is a key driver of fitness, directly influencing genetic contribution to future generations (Wadgymar et al., 2024), and is heavily influenced by nutrition (Smykal & Raikhel, 2015). In our study, flies fed with a protein-rich developmental diet produced significantly more viable offspring, consistent with previous research showing that higher yeast content in the developmental diet enhances reproductive performance in *Drosophila melanogaster* (Klepsatel et al., 2020) Similarly, protein rich adult diet also boosted fertility, consistent with previous findings (Lee et al., 2008; Schultzhaus & Carney, 2017). Protein-rich diets at both stages independently enhanced offspring production compared to carbohydrate-rich diets, with no significant interaction between developmental and adult diets. This aligns with previous work showing stage-specific dietary effects on fertility act independently (Ruchitha et al., 2024). The fertility boost from protein-rich diets is likely due to increased development of post-vitellogenic egg chambers in females (Ruchitha et al., 2024) due to abundant supply the dietary amino acids required for egg production (Alves et al., 2022; Grandison et al., 2009). Interestingly, while protein-rich developmental diets increase ovariole numbers, protein-rich adult diets enhance fecundity independently of ovariole number, highlighting distinct stage-specific mechanisms influencing reproductive output (Alves et al., 2022; Ruchitha et al., 2024).

Lifespan is another fundamental fitness trait and is reported to be majorly influenced by dietary macronutrient compositions (Lee et al., 2008; Mair et al., 2005). Building on past findings, our study demonstrates that developmental and adult diets impact male and female lifespans differently, highlighting the sexually dimorphic and stage-specific effects of macronutrient composition on longevity. Carbohydrate-rich developmental diets extended male lifespan, while female lifespan remained unaffected. The adult lifespan extension observed under low-protein, carbohydrate-rich developmental diets can be attributed to previously reported mechanisms, such as upregulation of the *Drosophila* transcription factor FOXO (dFOXO) in adult males (Gao et al., 2023) and reduced accumulation of harmful autotoxins under protein-restricted conditions (Stefana et al., 2017). Notably, both previous studies (Gao et al., 2023; Stefana et al., 2017 ) and ours consistently observed this developmental diet–induced lifespan extension exclusively in males, highlighting a robust, sex-specific response. While earlier studies reduced protein by keeping carbohydrate levels fixed – thereby lowering both caloric content and protein-to-carbohydrate ratio – our use of isocaloric diets isolates the effect of macronutrient composition and still yields lifespan extension. Together, these findings suggest that a lower protein-to-carbohydrate ratio in the developmental diet is sufficient to enhance male lifespan, independent of protein, carbohydrate, or caloric content alone. This provides a strong foundation for further exploration of the molecular and physiological mechanisms underlying this sexually dimorphic effect of developmental diet on lifespan. In contrast, the adult diet had a strong influence on lifespan with carbohydrate-rich diets significantly reducing lifespan compared to protein-rich diets in both sexes. While this aligns with the documented detrimental effect of excessive sucrose in adult diets (Lushchak et al., 2014), it contrasts with studies linking maximum lifespan to intermediate (Zanco et al., 2021) or lower (Jensen et al., 2015) protein-to-carbohydrate ratios. These discrepancies point to the complexity of dietary effects on lifespan, potentially arising from experimental differences across studies, including variations in the range of protein-to-carbohydrate ratios, caloric content, dietary formulations, or the genetic backgrounds of the flies. Notably, we observed no interaction between developmental and adult diets on lifespan in either sex, suggesting that dietary effects at each stage are independent. These results highlight the intricate interplay between dietary composition, life stage, sex, and their collective impact on lifespan. This interplay likely stems from complex energy allocation pathways, which also shape stress resistance traits such as starvation and desiccation resistance.

Survival success in the absence of food and water was influenced by the composition of diet consumed prior to these stresses, with notable differences between starvation and desiccation resistance. For starvation resistance, we observed a marginally significant effect of adult diet in females, with carbohydrate-rich diets enhancing survival compared to protein-rich diets.

This finding aligns with previous reports showing that lower protein-to-carbohydrate ratios improve starvation resistance ( Lee & Jang, 2014). The likely mechanism involves increased accumulation of lipid reserves under carbohydrate-rich conditions, which serve as key energy stores during food deprivation (Asiimwe et al., 2023). In contrast, male starvation resistance exhibited a more complex pattern, with a marginally significant interaction between developmental and adult diets, suggesting that the influence of adult nutrition may be modulated by the larval dietary environment in a sex-specific manner. This highlights the potential for context-dependent physiological responses in males and warrants further investigation into the developmental programming of resource storage, allocation and utilization in adults. For desiccation resistance, adult diet again played a dominant role, with carbohydrate-rich diets consistently improving survival under desiccation stress compared to protein-rich diets in both sexes. This effect is likely mediated by enhanced glycogen accumulation (Djawdan et al., 1998), as glycogen serves both as a metabolic energy source and a water-binding molecule that improves hydration status during desiccation (Archer et al., 2007; Gibbs A & Rose, 2004). In contrast to starvation resistance, there was no evidence of interaction between developmental and adult diets for desiccation resistance in either sex, nor any main effect of developmental diet, suggesting that this trait is more directly shaped by the immediate nutritional environment during adulthood. Together, these findings underscore the predominant role of adult diet in shaping stress resistance traits, while also demonstrating that the interplay between developmental and adult nutrition can vary in a trait- and sex-specific manner. Future studies quantifying lipid, glycogen, and water content across life stages and diets would help clarify the physiological mechanisms underlying these patterns.

Below, we highlight four overarching insights from our study and their broader implications. First, there is no universally best diet regime across the traits and life stages examined in our study. The effects of dietary composition on traits vary by life stages and the specific trait in question, emphasizing the importance of timing and context in dietary interventions. While our findings are based on two contrasting P:C ratios (0.25 and 0.7), they align with previous studies showing non-monotonic responses across broader P:C ranges (Lee, 2015; Zanco et al., 2021). This complexity underscores the trade-offs organisms must navigate to maximize fitness and has important implications for how populations adapt to environmental changes, such as habitat loss or climate change, which influence the availability and quality of food resources.

Second, our study highlights that, despite undergoing complete metamorphosis, during which larval tissues are largely replaced, developmental nutrition in *Drosophila* still plays a vital role in shaping adult traits. Previous research suggests that larval fat cells, acting as nutrient reservoirs during early adult stage (Aguila et al., 2007; Andersen et al., 2010), may drive some of the effects of developmental diet, but further research is needed to fully understand the mechanisms involved. The influence of early nutrition emphasizes its long-term impact on fitness and adaptability across diverse ecological contexts, with broader implications for evolutionary biology.

Third, our full factorial experimental design allowed us to examine potential interactions between developmental and adult diets. While we found no significant interactions for most of the five measured life-history traits, indicating largely additive effects of stage-specific nutrition, a marginal interaction for male starvation resistance suggests that developmental and adult diets can interact in a trait- and sex-specific manner. This observation highlights the need for further investigation into the mechanisms through which stage-specific diets influence life-history traits, particularly in a sex-dependent context.

Finally, by evaluating pairwise associations between dietary effects on life-history traits, we found a context-dependent associations of dietary effects on these traits, varying with life stage and sex. Developmental diet showed no significant associations with the measured adult traits, whereas adult dietary effects showed significant and sex-specific patterns. In both males and females, adult dietary effects on fertility and longevity were positively associated, suggesting that diets promoting higher reproductive output also enhance survival under adult dietary conditions. This observation is interesting given the canonical fecundity-longevity trade-off, often attributed to resource constraints (Hunt et al., 2004; Partridge et al., 2005) and may have important evolutionary implications. In contrast, adult dietary effects on desiccation resistance were negatively associated with their effects on both fertility and longevity in both sexes. In females, starvation resistance also showed negative associations with fertility and longevity, reflecting resource allocation trade-offs. Notably, starvation resistance was positively associated with desiccation resistance in females, suggesting shared physiological mechanisms for stress tolerance. These associations were absent in males, highlighting potential sex-specific differences in resource allocation strategies. The observed negative associations between dietary effects on fertility and stress resistance traits align with their established trade-offs (Wang et al., 2001) However, the negative association between dietary effects on longevity and stress resistance traits is counterintuitive, as stress resistance is often used as a proxy for lifespan in some species (Lithgow & Walker, 2002). These findings caution against extrapolating evolutionary consequences or using stress resistance as a proxy for lifespan without considering dietary contexts. Overall, these context-dependent associations allude to differential resource allocation strategies and phenotypic plasticity in males and females depending on the nutritional environments across life stages.

## 5. Conclusion

Our study provides a systematic examination of how dietary macronutrient composition during developmental and adult stages influences multiple life-history traits in *Drosophila melanogaster*, demonstrating stage- and sex-specific patterns. Developmental diet played a key role in shaping adult body size and lifespan in males, highlighting an enduring impact of early nutrition. Additionally, adult diet significantly influenced traits such as fertility, lifespan, and stress resistance, with carbohydrate-rich diets enhancing survival under starvation and desiccation stress but reducing fertility and longevity. These results underscore the complex interplay between dietary composition, life stage, and sex in driving resource allocation strategies and fitness outcomes. Furthermore, the analysis of context-dependent associations between dietary effects showed both expected trade-offs, such as the negative associations between stress resistance and reproductive or longevity traits, and counterintuitive findings, such as the positive association between fertility and longevity under adult dietary conditions. The sexually dimorphic nature of these associations emphasizes the need for sex-specific considerations in evolutionary models. Future research should focus on understanding the molecular and physiological mechanisms underlying these patterns, including nutrient-sensing pathways and the role of metabolic reserves such as lipids and glycogen. Additionally, exploring genetic adaptations under contrasting nutritional environments could provide valuable insights into how populations would respond to changing ecological conditions. These findings have far-reaching implications for understanding how nutrition drives phenotypic plasticity and fitness trade-offs, shaping ecological dynamics and evolutionary trajectories in populations.

## Author contributions

Conceptualization: Mohankumar Chandrakanth, Sudipta Tung

Design of the experiment: Mohankumar Chandrakanth, Nishant Kumar, Chand Sura, Sudipta Tung

Data curation: Mohankumar Chandrakanth, Nishant Kumar, Chand Sura, Sudipta Tung

Formal analysis: Mohankumar Chandrakanth

Funding acquisition: Sudipta Tung

Supervision: Sudipta Tung

Visualization: Mohankumar Chandrakanth

Writing – original draft: Mohankumar Chandrakanth, Sudipta Tung

Writing – review & editing: Mohankumar Chandrakanth, Sudipta Tung

## Acknowledgements

The authors thank Prof. Sutirth Dey from IISER Pune for helping with the statistical analysis of this work. We also thank Mr. Devashish Kumar from Ashoka University for his valuable comments on the manuscript. MC acknowledges the support from the Department of Biotechnology, Govt. of India through Junior Research Fellowship. NK and CS acknowledge the support of the Research and Development Office, Ashoka University. ST acknowledges the support of DBT/Wellcome Trust India Alliance Early Career Fellowship (#IA/E/18/1/504347) and Ashoka University.

## Data availability

Data used in this manuscript has been uploaded on Dryad repository and can be accessed using the link below:

http://datadryad.org/share/IxTYtbpR1beH5Krj6YPNjeknRYDh9ZCickN0LZ3kP_U

