## Supplementary for "Context-dependent effects of developmental and adult diet on life-history traits in *Drosophila melanogaster*"

**Text S1. Egg collection protocol**

To collect eggs from flies housed in plexiglass cages, we used an egg-laying substrate referred to as a cut-plate—a food plate with cuts to create vertical surfaces for egg deposition—prepared with the corresponding adult diet. Flies were allowed to lay eggs on the cut-plate overnight for approximately 14 hours. During collection, the cut-plate was carefully removed from each cage, and the vertical surfaces were rinsed with a small amount of distilled water to loosen the eggs. Autoclaved 2 mL microcentrifuge tubes (MCTs) were prepared, one for each cage, containing 750 µL of distilled water. Using a fine, wet paintbrush (size 000), eggs were gently scraped from the cut-plate and carefully transferred into the corresponding MCT until the volume of eggs reached 500 µL mark. To remove food particles attached to the eggs, 750 µL of distilled water was added to each tube. The tubes were gently tapped to break up clumps and slowly inverted three times to clean the eggs. After allowing the eggs to settle, 750 µL of the supernatant containing food debris was carefully removed and discarded. Finally, 50 µL of well-mixed egg suspension was aliquoted into culture bottles containing 50 mL of fresh food. Based on prior standardization, this corresponded to an inoculum of approximately 300-350 eggs per bottle. The bottles were incubated at 25°C under constant light and >60% humidity.

**Table S1. Composition of isocaloric experimental diets**

| **P:C ratio** | **0.25**  (Carbohydrate-rich diet) | **0.4**  (Baseline diet) | **0.7**  (Protein-rich diet) |
| --- | --- | --- | --- |
| Water (mL) | 1000 | 1000 | 1000 |
| Corn (g) | 50 | 50 | 50 |
| Sugar (g) | 79.7 | 50 | 10.3 |
| Yeast (g) | 67.6 | 100 | 143.5 |
| Agar (g) | 15 | 15 | 15 |
| Extra water (mL) | 200 | 200 | 200 |
| Benzoate (mL) | 40 | 40 | 40 |
| Propionic acid (mL) | 20 | 20 | 20 |

The flies derived carbohydrates from sugar (3.87 KCal/g) and corn (3.33 KCal/g) and both proteins and carbohydrates from yeast (3.54 KCal/g). The protein-to-carbohydrate (P:C) ratio was calculated based on the total protein and carbohydrate content of each dietary ingredient. Per gram, yeast contains 0.48 g of protein and 0.37 g of carbohydrates, while corn contains 0.07 g of protein and 0.78 g of carbohydrates (values based on package nutritional information). The total energy content was standardized to 714 kcal per litre of food across all diet regimes.

**Table S2. Results of full factorial analysis of wing length, lifespan, starvation resistance, and desiccation resistance**

Wing length (model: glm(Wing_length ~ Larval_food*Adult_food*Sex, family = "gaussian", data))

Sum Sq Df F values Pr(>F)

Larval_food	0.00329	1	3.8546 0.0508 .

Adult_food	0.00033	1	0.3840 0.5361

Sex	0.45054	1 528.1825 <2e-16 ***

Larval_food:Adult_food	0.00161	1	1.8886 0.1707

Larval_food:Sex	0.00006	1	0.0753 0.7841

Adult_food:Sex	0.00064	1	0.7500 0.3874

Larval_food:Adult_food:Sex 0.00125	1	1.4649 0.2274

Residuals	0.19790 232

Lifespan (model:coxme(Surv(Time, Status)~Larval_food*Adult_food*Sex +(1|Vial), data = data))

Df	Chisq Pr(>Chisq)

Larval_food	1 0.1324	0.715982

Adult_food	1 45.7918 1.315e-11 ***

Sex	1 79.2483 < 2.2e-16 ***

Larval_food:Adult_food	1 3.9837	0.045942 *

Larval_food:Sex	1 3.9490	0.046899 *

Adult_food:Sex	1 9.9800	0.001583 **

Larval_food:Adult_food:Sex 1 6.6489	0.009922 **

Starvation resistance (model:coxme(Surv(Time, Status)~Larval_food*Adult_food*Sex +(1|Vial), data = data))

Df	Chisq Pr(>Chisq)

Larval_food	1 0.6225	0.43012

Adult_food	1 3.2651	0.07077 .

Sex	1 66.1157	4.252e-16 ***

Larval_food:Adult_food	1 0.4012	0.52645

Larval_food:Sex	1 0.1514	0.69720

Adult_food:Sex	1 0.5391	0.46279

Larval_food:Adult_food:Sex 1 5.2672	0.02173 *

Desiccation resistance (model:coxme(Surv(Time, Status)~Larval_food*Adult_food*Sex +(1|Vial), data = data))

Df	Chisq Pr(>Chisq)

Larval_food	1	0.5436	0.460941

Adult_food	1	6.9744	0.008269 **

Sex	1 159.3538 < 2.2e-16 ***

Larval_food:Adult_food	1	0.0288	0.865195

Larval_food:Sex	1	0.9935	0.318882

Adult_food:Sex	1	0.4055	0.524249

Larval_food:Adult_food:Sex 1	0.0399	0.841613

**Table S3: The developmental diet and adult diet impact on wing length in females and male *Drosophila*.**

| **Wing length** | | | | | | |
| --- | --- | --- | --- | --- | --- | --- |
| **Females** | | | | | | |
|  | **Sum of Squares** | **Degree of freedom** | **F Value** | **p value** | **Slope (β) with C diet as reference** | **95% CI** |
| Developmental diet | 0.001677 | 1 | 1.9000 | 0.1707 | -0.0148 | (-0.0298, 0.0002) |
| Adult diet | 0.000211 | 1 | 0.2396 | 0.6254 | -0.00467 | (-0.0197, 0.0104) |
| Developmental diet × Adult food | 0.001611 | 1 | 1.8248 | 0.1794 | 0.0147 | (-0.0066, 0.0359) |
| **Males** | | | | | | |
| Developmental diet | 0.005613 | 1 | 6.8190 | 0.01021 | -0.0119 | (-0.0264, 0.0026) |
| Adult diet | 0.000229 | 1 | 0.2784 | 0.59875 | 0.00456 | (-0.0099, 0.0191) |
| Developmental diet × Adult food | 0.000097 | 1 | 0.1180 | 0.73187 | -0.0036 | (-0.0241, 0.0169) |

**Table S4: The effects of developmental diet and adult diet on fertility in *Drosophila*.**

| **Fertility** | | | | | | |
| --- | --- | --- | --- | --- | --- | --- |
|  | **Sum of Squares** | **Degree of freedom** | **F Value** | **p value** | **Slope (β) with C diet as reference** | **95% CI** |
| Developmental diet | 4.302 | 1 | 4.3844 | 0.03845 | 0.137 | (-0.0061, 0.281) |
| Adult diet | 60.347 | 1 | 61.5082 | 2.377e-12 | 0.417 | (0.278,  0.556) |
| Developmental diet × Adult food | 0.419 | 1 | 0.4266 | 0.51494 | -0.064 | (-0.258, 0.130) |

**Table S5: The effects of developmental diet and adult diet on lifespan in females and male *Drosophila*.**

| **Longevity** | | | | | |
| --- | --- | --- | --- | --- | --- |
| **Females** | | | | | |
|  | **Degree of freedom** | **χ^2^** | **p value** | **Slope (β) with C diet as reference** | **95% CI** |
| Developmental diet | 1 | 0.1045 | 0.7465 | -0.052 | (-0.3691, 0.2646) |
| Adult diet | 1 | 35.7503 | 2.243e-09 | -1.03 | (-1.368, -0.6926) |
| Developmental diet × Adult food | 1 | 2.4531 | 0.1173 | 0.371 | (0.3716,  0.8367) |
| **Males** | | | | | |
| Developmental diet | 1 | 6.2015 | 0.01276 | 0.406 | (0.0864,  0.7256) |
| Adult diet | 1 | 6.5062 | 0.01075 | -0.416 | (-0.7366,  -0.0964) |
| Developmental diet × Adult food | 1 | 2.8461 | 0.09160 | -0.387 | (-0.8381,  0.0627) |

**Table S6: The effects of developmental diet and adult diet on starvation resistance in females and male *Drosophila*.**

| **Starvation resistance** | | | | | |
| --- | --- | --- | --- | --- | --- |
| **Females** | | | | | |
|  | **Degree of freedom** | **χ^2^** | **p value** | **Slope (β) with C diet as reference** | **95% CI** |
| Developmental diet | 1 | 0.9126 | 0.33942 | -0.15 | (-0.4607, 0.1587) |
| Adult diet | 1 | 3.9910 | 0.04574 | 0.316 | (0.0059,  0.6269) |
| Developmental diet × Adult food | 1 | 0.5053 | 0.47719 | 0.159 | (-0.2803, 0.5993) |
| **Males** | | | | | |
| Developmental diet | 1 | 2.1311 | 0.14433 | -0.235 | (-0.5509, 0.0805) |
| Adult diet | 1 | 4.5106 | 0.03368 | 0.342 | (0.0263, 0.6577) |
| Developmental diet × Adult food | 1 | 3.6873 | 0.05483 | -0.435 | (-0.8797, 0.009) |

**Table S7: The effects of developmental diet and adult diet on desiccation resistance in females and male *Drosophila*.**

| **Desiccation resistance** | | | | | |
| --- | --- | --- | --- | --- | --- |
| **Females** | | | | | |
|  | **Degree of freedom** | **χ^2^** | **p value** | **Slope (β) with C diet as reference** | **95% CI** |
| Developmental diet | 1 | 0.5436 | 0.460940 | -0.119 | (-0.4356,  0.1975) |
| Adult diet | 1 | 7.3242 | 0.006803 | 0.435 | (0.1201, 0.7511) |
| Developmental diet × Adult food | 1 | 0.0182 | 0.892633 | -0.03 | (-0.4764,  0.415) |
| **Males** | | | | | |
| Developmental diet | 1 | 2.9933 | 0.0836101 | -0.279 | (-0.5965,  0.0371) |
| Adult diet | 1 | 11.6671 | 0.0006361 | 0.544 | (0.2321, 0.8574) |
| Developmental diet × Adult food | 1 | <0.0001 | 0.9977910 | 0.0006 | (-0.4423, 0.4435) |

**Table S8. Model fit metrics (R^2^) for each trait and sex**

| **Trait** | **Sex** | **Model Fit Metric** | **Value** |
| --- | --- | --- | --- |
| Wing length | Male | Adjusted R^2^ | 0.059 |
|  | Female | Adjusted R^2^ | 0.033 |
| Fertility | - | Nagelkerke’s R^2^ | 0.526 |
| Longevity | Male | Nagelkerke’s R^2^ | 0.094 |
|  | Female | Nagelkerke’s R^2^ | 0.115 |
| Starvation resistance | Male | Nagelkerke’s R^2^ | 0.055 |
|  | Female | Nagelkerke’s R^2^ | 0.04 |
| Desiccation resistance | Male | Nagelkerke’s R^2^ | 0.083 |
|  | Female | Nagelkerke’s R^2^ | 0.045 |

| **Table S9. Descriptive statistics and effect size for all traits across the sexes** | | | | | | | | | |
| --- | --- | --- | --- | --- | --- | --- | --- | --- | --- |
| **Female** | | | | | | | | | |
| Traits | Stage | Carbohydrate-rich diet | | | Protein-rich diet | | | Cohen's *d* | Inference |
|  |  | Sample | Mean | SE | Sample | Mean | SE |  |  |
| Wing Length | Larva | 60 | 1.4 | 0.00341 | 60 | 1.39 | 0.00422 | 0.34 | Small |
|  | Adult | 60 | 1.4 | 0.00419 | 60 | 1.4 | 0.00351 | 0 | Very small |
| Fertility | Larva | 60 | 28.2 | 1.16 | 60 | 31.2 | 1.32 | 0.31 | Small |
|  | Adult | 60 | 24.1 | 0.94 | 60 | 35.3 | 1.11 | 1.41 | Large |
| Longevity | Larva | 158 | 32.9 | 0.972 | 155 | 32.1 | 0.893 | 0.07 | Very small |
|  | Adult | 158 | 27.8 | 0.882 | 155 | 37.3 | 0.821 | 0.89 | Large |
| Starvation Resistance | Larva | 161 | 69.6 | 0.94 | 159 | 70.1 | 1.32 | 0.03 | Very small |
|  | Adult | 161 | 72.6 | 1.02 | 159 | 67 | 1.22 | 0.39 | Small |
| Desiccation Resistance | Larva | 160 | 30.6 | 0.392 | 159 | 30.8 | 0.486 | 0.04 | Very small |
|  | Adult | 160 | 31.8 | 0.454 | 159 | 29.6 | 0.41 | 0.4 | Small |
| **Male** | | | | | | | | | |
| Traits | Stage | Carbohydrate-rich | | | Protein-rich | | | Cohen's *d* | Inference |
|  |  | Sample | Mean | SE | Sample | Mean | SE |  |  |
| Wing Length | Larva | 60 | 1.23 | 0.00379 | 60 | 1.22 | 0.00357 | 0.35 | Small |
|  | Adult | 60 | 1.22 | 0.00395 | 60 | 1.23 | 0.0036 | 0.34 | Small |
| Fertility | Larva | 60 | 28.2 | 1.16 | 60 | 31.2 | 1.32 | 0.31 | Small |
|  | Adult | 60 | 24.1 | 0.94 | 60 | 35.3 | 1.11 | 1.41 | Large |
| Longevity | Larva | 160 | 43.8 | 0.965 | 158 | 41.3 | 1.03 | 0.20 | Small |
|  | Adult | 158 | 39.4 | 0.962 | 160 | 45.7 | 0.981 | 0.51 | Medium |
| Starvation Resistance | Larva | 159 | 54.8 | 0.91 | 159 | 58.5 | 0.902 | 0.32 | Small |
|  | Adult | 159 | 57.2 | 0.937 | 159 | 56.1 | 0.897 | 0.10 | Very small |
| Desiccation Resistance | Larva | 160 | 20.8 | 0.263 | 159 | 21.5 | 0.326 | 0.19 | Very small |
|  | Adult | 159 | 22.2 | 0.297 | 160 | 20.1 | 0.272 | 0.58 | Medium |
| *Note: Same fertility data is written in both female and male table. | | | | | | | | | |

**Table S10. Dietary-effect indices between the effect of developmental diet on adult life-history traits**

| Trait combination | Male | | | | Female | | | |
| --- | --- | --- | --- | --- | --- | --- | --- | --- |
|  | **DEI*_ij_***  $\frac{\mathrm{lnRR}_{\mathrm{trait}i}}{\mathrm{lnRR}_{\mathrm{trait}j}}$ | **SE*_ij_***    $\sqrt{\left( \frac{V_{\mathrm{trait}i}}{{\mathrm{lnRR}_{\mathrm{trait}j}}^{2}}+\frac{{\mathrm{lnRR}_{\mathrm{trait}i}}^{2}V_{\mathrm{trait}j}}{{\mathrm{lnRR}_{\mathrm{trait}j}}^{4}} \right)}$ | **Z**  $\frac{\mathrm{TAI}_{ij}}{\mathrm{SE}_{ij}}$ | ***p***  **value** | **DEI*_ij_***  $\frac{\mathrm{lnRR}_{\mathrm{trait}i}}{\mathrm{lnRR}_{\mathrm{trait}j}}$ | **SE*_ij_***    $\sqrt{\left( \frac{V_{\mathrm{trait}i}}{{\mathrm{lnRR}_{\mathrm{trait}j}}^{2}}+\frac{{\mathrm{lnRR}_{\mathrm{trait}i}}^{2}V_{\mathrm{trait}j}}{{\mathrm{lnRR}_{\mathrm{trait}j}}^{4}} \right)}$ | **Z**  $\frac{\mathrm{TAI}_{ij}}{\mathrm{SE}_{ij}}$ | ***p***  **value** |
| Wing length vs Fertility | -0.08 | 0.06 | -1.28 | 0.20 | -0.07 | 0.06 | -1.25 | 0.21 |
| Wing length vs Longevity | 0.14 | 0.11 | 1.30 | 0.19 | 0.29 | 0.50 | 0.58 | 0.56 |
| Wing length vs Starvation resistance | -0.12 | 0.08 | -1.60 | 0.11 | -1.00 | 3.29 | -0.30 | 0.76 |
| Wing length vs Desiccation resistance | -0.25 | 0.20 | -1.26 | 0.21 | -1.10 | 3.49 | -0.32 | 0.75 |
| Fertility vs Longevity | -1.72 | 1.40 | -1.23 | 0.22 | -4.11 | 7.18 | -0.57 | 0.57 |
| Fertility vs Starvation resistance | 196.00 | 8630.56 | 0.02 | 0.98 | 14.12 | 46.53 | 0.30 | 0.76 |
| Fertility vs Desiccation resistance | 3.05 | 2.56 | 1.20 | 0.23 | 15.52 | 49.28 | 0.31 | 0.75 |
| Longevity vs Starvation resistance | -113.94 | 5017.27 | -0.02 | 0.98 | -3.44 | 12.50 | -0.28 | 0.78 |
| Longevity vs Desiccation resistance | -1.78 | 1.46 | -1.21 | 0.22 | -3.78 | 13.33 | -0.28 | 0.78 |
| Starvation resistance vs Desiccation resistance | 1.97 | 1.36 | 1.45 | 0.15 | 1.10 | 4.94 | 0.22 | 0.82 |

**Table S11. Dietary-effect indices between the effect of adult diet on adult life-history traits**

| Trait combination | Male | | | | Female | | | |
| --- | --- | --- | --- | --- | --- | --- | --- | --- |
|  | **DEI*_ij_***  $\frac{\mathrm{lnRR}_{\mathrm{trait}i}}{\mathrm{lnRR}_{\mathrm{trait}j}}$ | **SE*_ij_***    $\sqrt{\left( \frac{V_{\mathrm{trait}i}}{{\mathrm{lnRR}_{\mathrm{trait}j}}^{2}}+\frac{{\mathrm{lnRR}_{\mathrm{trait}i}}^{2}V_{\mathrm{trait}j}}{{\mathrm{lnRR}_{\mathrm{trait}j}}^{4}} \right)}$ | **Z**  $\frac{\mathrm{TAI}_{ij}}{\mathrm{SE}_{ij}}$ | ***p***  **value** | **DEI*_ij_***  $\frac{\mathrm{lnRR}_{\mathrm{trait}i}}{\mathrm{lnRR}_{\mathrm{trait}j}}$ | **SE*_ij_***    $\sqrt{\left( \frac{V_{\mathrm{trait}i}}{{\mathrm{lnRR}_{\mathrm{trait}j}}^{2}}+\frac{{\mathrm{lnRR}_{\mathrm{trait}i}}^{2}V_{\mathrm{trait}j}}{{\mathrm{lnRR}_{\mathrm{trait}j}}^{4}} \right)}$ | **Z**  $\frac{\mathrm{TAI}_{ij}}{\mathrm{SE}_{ij}}$ | ***p***  **value** |
| Wing length vs Fertility | 0.02 | 0.01 | 1.82 | 0.07 | 0 | 0.010 | 0 | 1 |
| Wing length vs Longevity | 0.06 | 0.03 | 1.73 | 0.08 | 0 | 0.013 | 0 | 1 |
| Wing length vs Starvation resistance | -0.42 | 0.54 | -0.77 | 0.44 | 0 | 0.049 | 0 | 1 |
| Wing length vs Desiccation resistance | -0.08 | 0.05 | -1.76 | 0.08 | 0 | 0.054 | 0 | 1 |
| Fertility vs Longevity | 2.57 | 0.66 | 3.92 | 0.0001 | 1.30 | 0.24 | 5.39 | 7.12e-08 |
| Fertility vs  Starvation resistance | -19.66 | 23.29 | -0.84 | 0.40 | -4.75 | 1.50 | -3.18 | 0.001 |
| Fertility vs Desiccation resistance | -3.84 | 0.89 | -4.31 | 0.00002 | -5.32 | 1.63 | -3.26 | 0.001 |
| Longevity vs Starvation resistance | -7.64 | 9.15 | -0.83 | 0.40 | -3.66 | 1.15 | -3.17 | 0.002 |
| Longevity vs Desiccation resistance | -1.49 | 0.43 | -3.43 | 0.0006 | -4.10 | 1.26 | -3.26 | 0.001 |
| Starvation resistance vs Desiccation resistance | 0.20 | 0.23 | 0.84 | 0.40 | 1.12 | 0.45 | 2.51 | 0.012 |

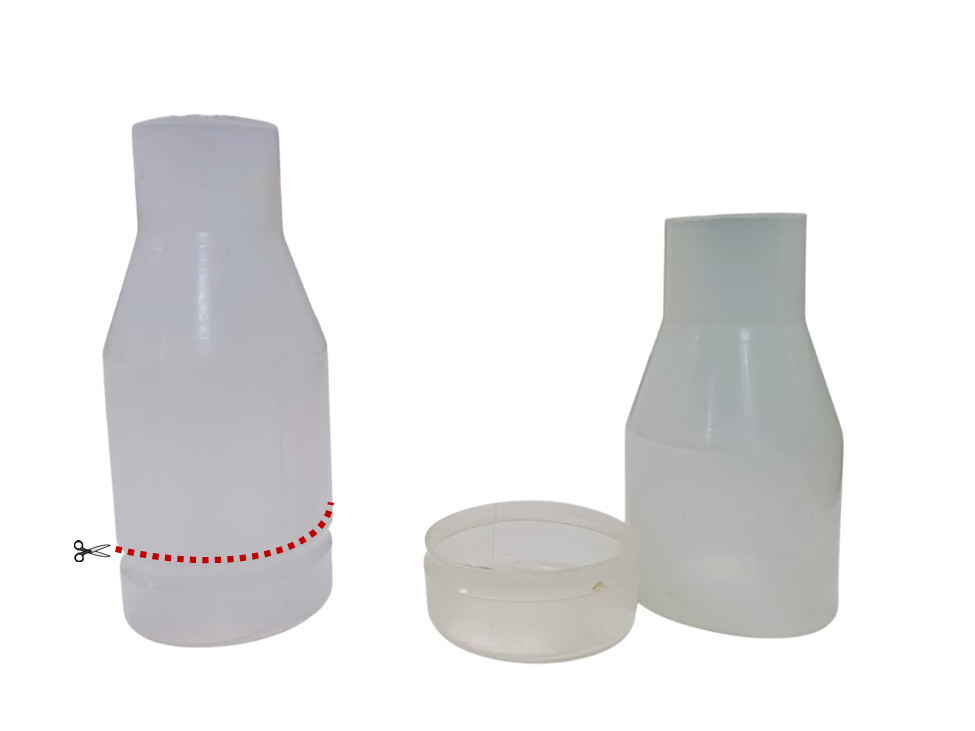

**Figure S1. Modified fly-culture bottle for transitioning from developmental to adult diet at the late pupal stage.** The intact plastic fly-culture bottle (left) is shown with a red dotted line indicating where it is cut to create two separate parts: a removable food cup and an upper container (right). This design allows easy swapping of the developmental diet with the adult diet.

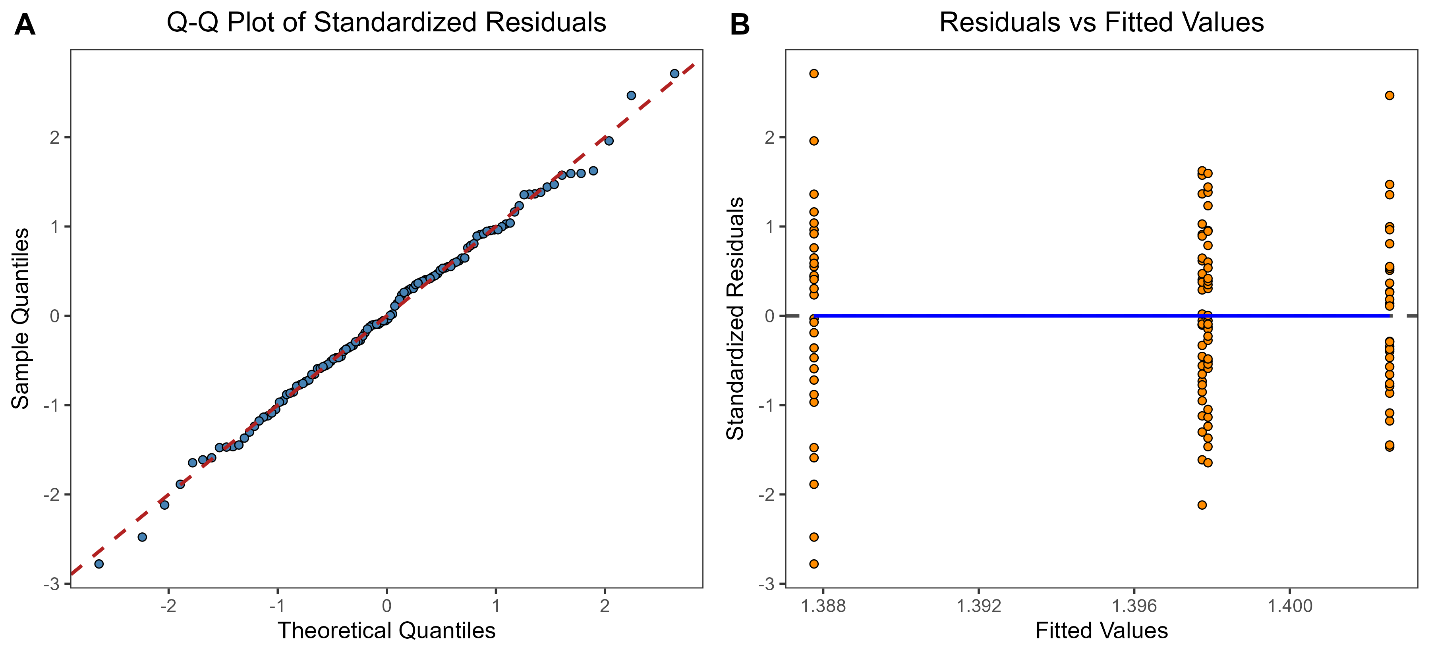

**Figure S2.** **Residual diagnostics for the wing length model in females.** **(A)** Q-Q plot of standardized residuals shows alignment with the theoretical normal distribution, indicating normality. **(B)** Residuals vs. fitted values plot shows no clear pattern or heteroscedasticity, supporting the assumption of homoscedasticity.

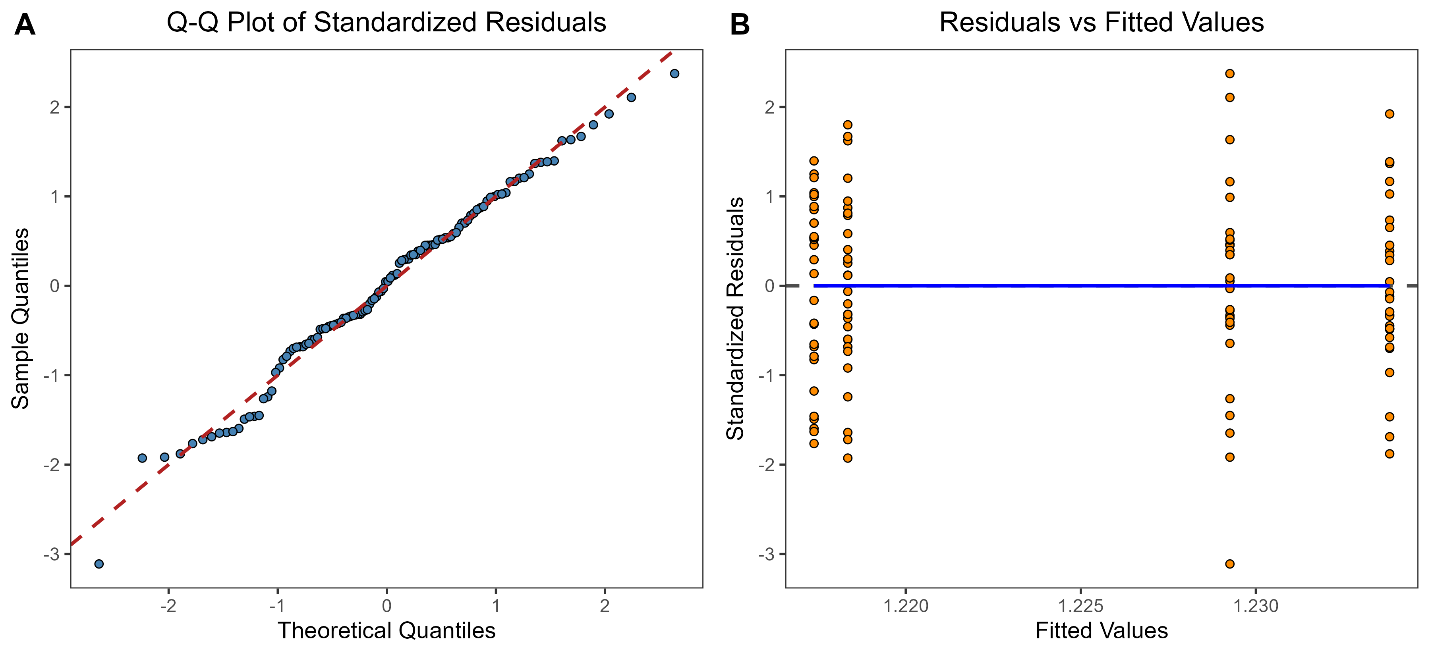

**Figure S3.** **Residual diagnostics for the wing length model in males.** (A) Q-Q plot of standardized residuals shows alignment with the theoretical normal distribution, indicating normality. (B) Residuals vs. fitted values plot shows no clear pattern or heteroscedasticity, supporting the assumption of homoscedasticity.

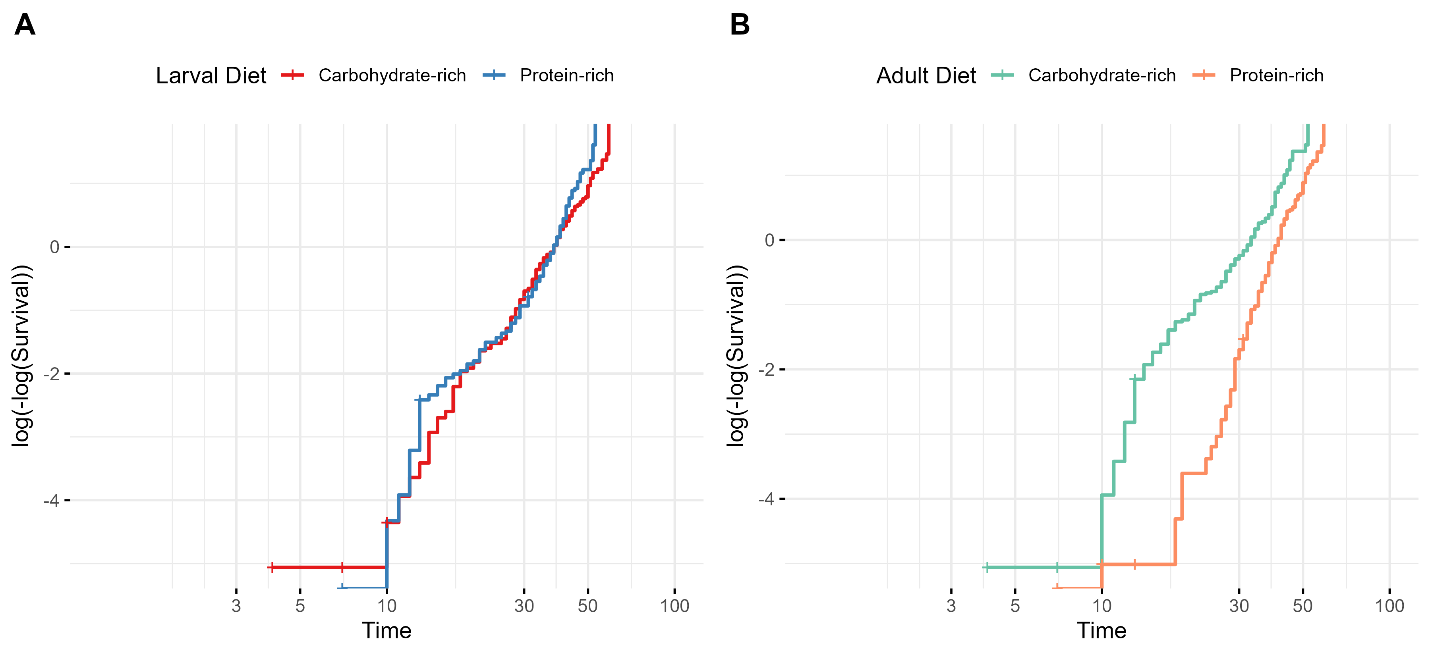

**Figure S4. Log-minus-log survival plots for female lifespan data stratified by larval diet (A) and adult diet (B).**

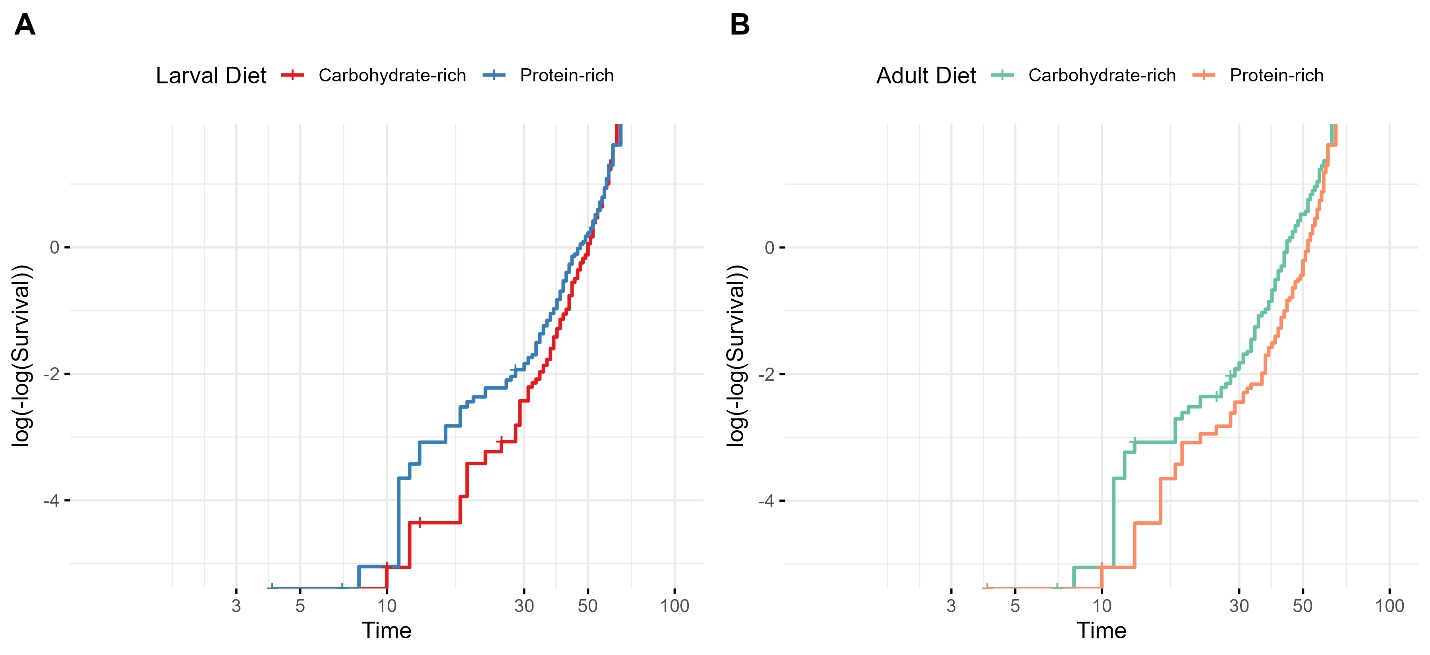

**Figure S5. Log-minus-log survival plots for male lifespan data stratified by larval diet (A) and adult diet (B).**

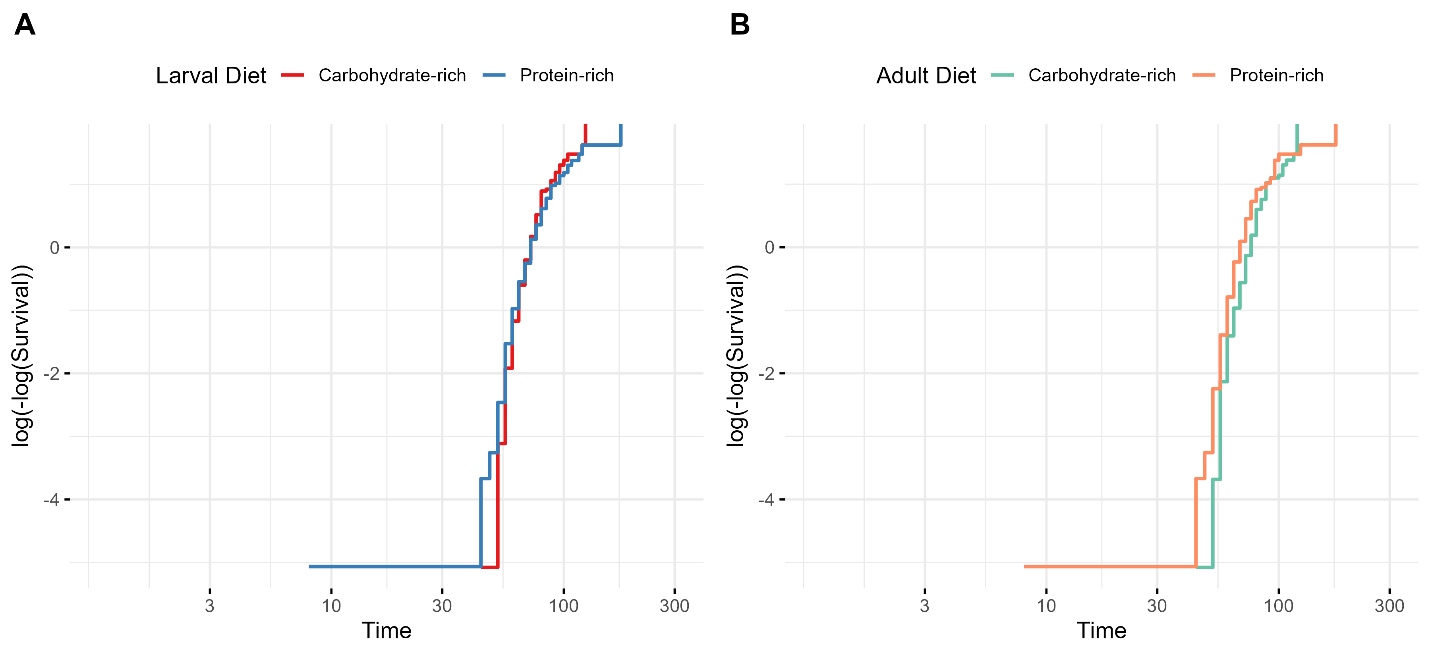

**Figure S6. Log-minus-log survival plots for female starvation resistance data stratified by larval diet (A) and adult diet (B).**

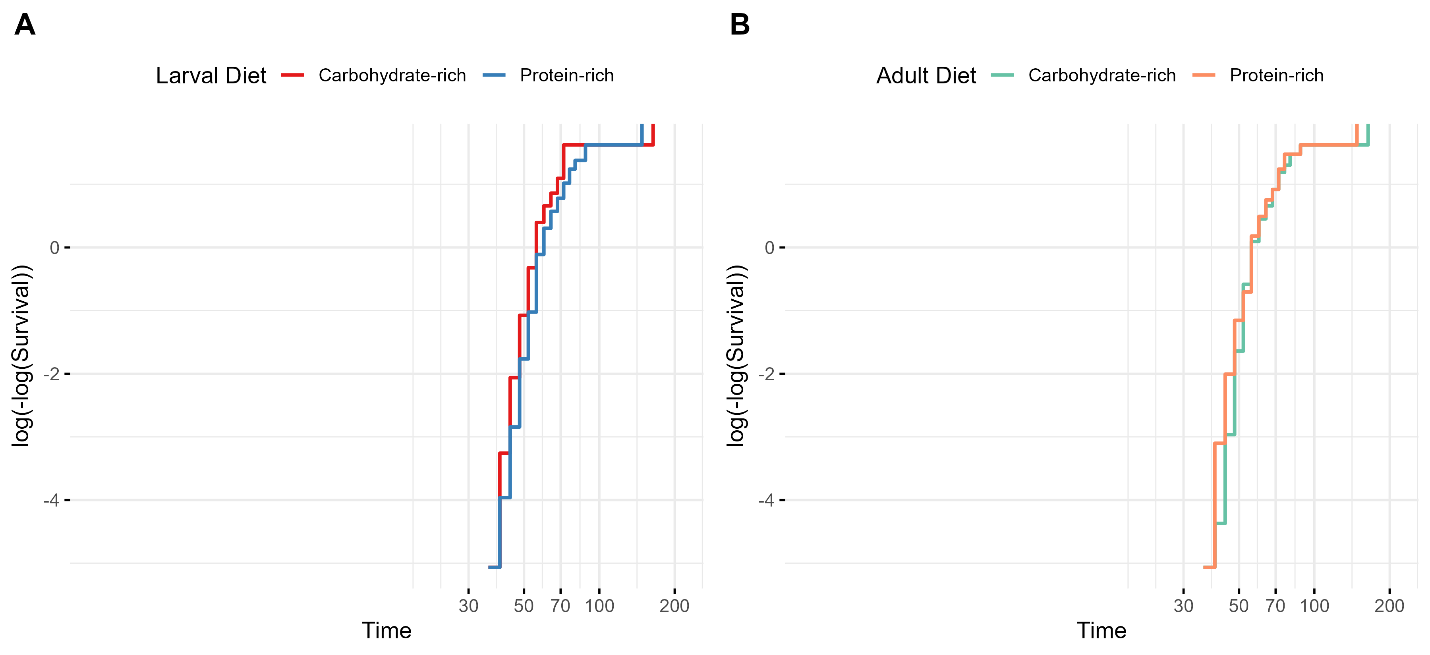

**Figure S7. Log-minus-log survival plots for male starvation resistance data stratified by larval diet (A) and adult diet (B).**

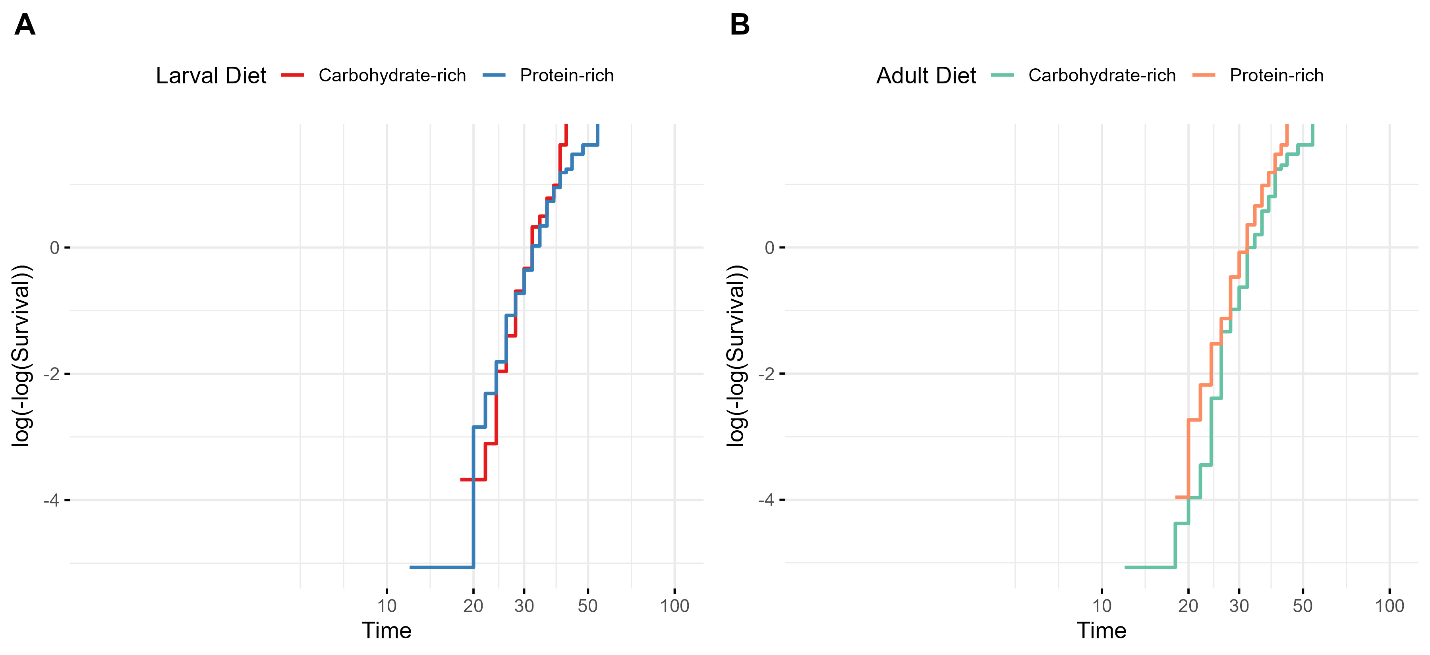

**Figure S8. Log-minus-log survival plots for female desiccation resistance data stratified by larval diet (A) and adult diet (B).**

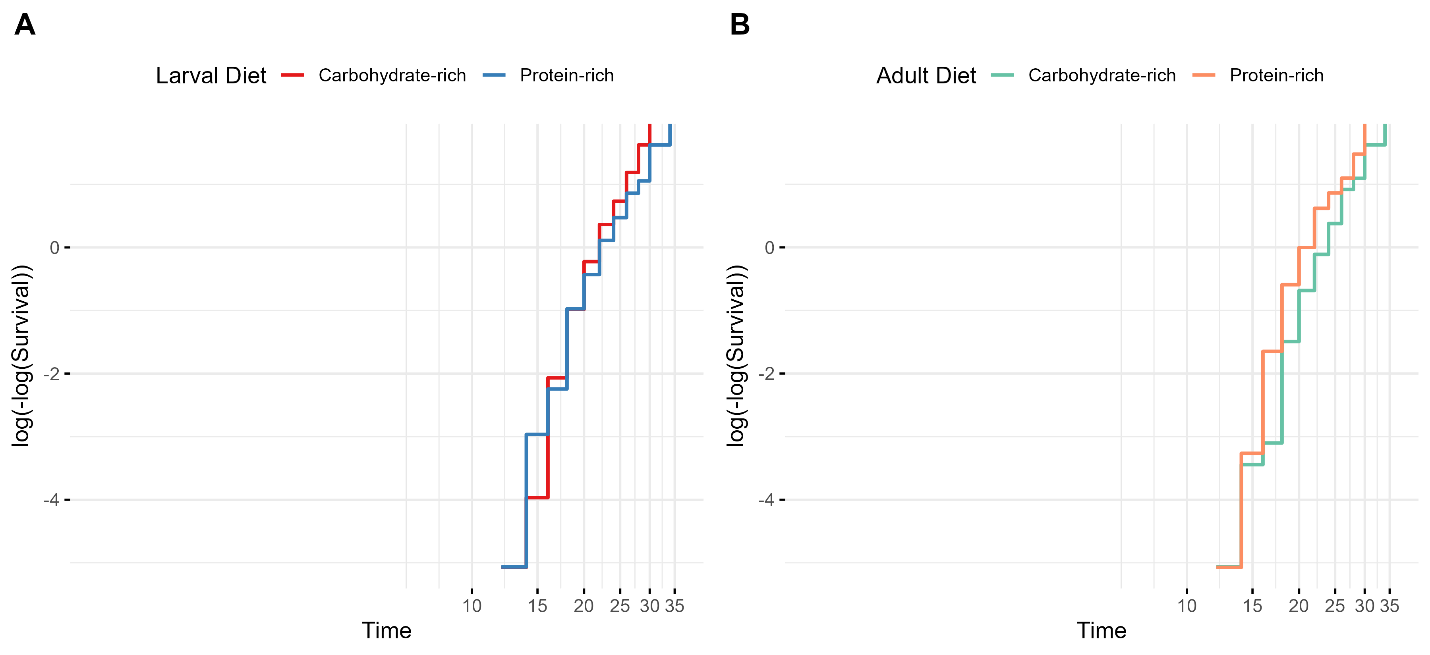

**Figure S9. Log-minus-log survival plots for male desiccation resistance data stratified by larval diet (A) and adult diet (B).**

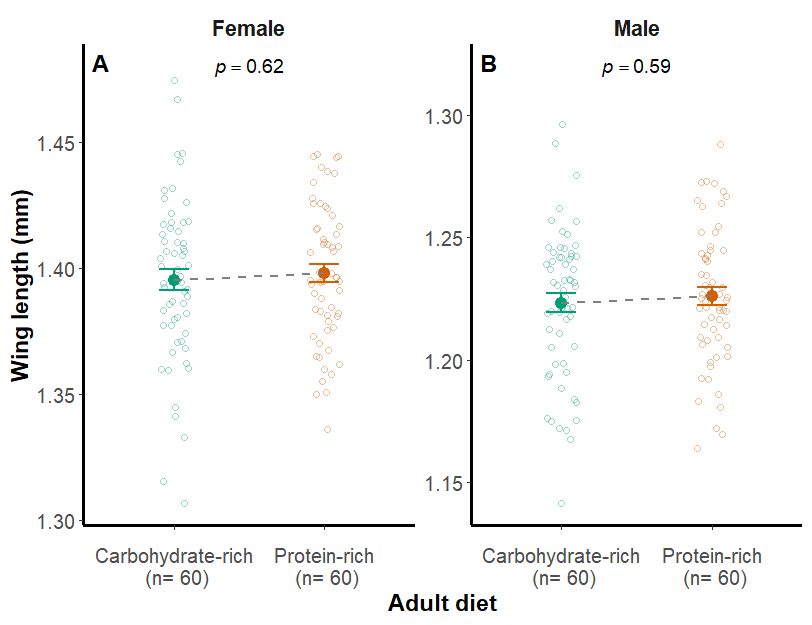

**Figure S10.** Effect of adult diet composition on wing length. The graphs depict the impact of carbohydrate-rich (C) and protein-rich (P) adult diet on adult wing length (a proxy for body size). Panels (A) and (B) show the effects of adult diet on wing length in females and males, respectively. Error bars represent the standard error of the mean (± SE) for each diet. The filled circles indicate mean wing length, while small open circles denote wing length of individual flies in each group. Teal green and orange-red color represents the carbohydrate-rich and protein-rich isocaloric diets, respectively. n denotes the sample size; p values represent statistical significance levels.
